# CRISPR/Cas9-based somatic knock-in of reporters in the avian embryo in ovo

**DOI:** 10.1101/2024.10.10.617291

**Authors:** Alciades Petit Vargas, Baptiste Mida, Rosette Goïame, Olinda Alegria-Prevot, Bojana Djelic, Evelyne Fischer, Samuel Tozer, Jérôme Gros, Marie Manceau, Xavier Morin

**Affiliations:** Institut de Biologie de l’ENS (IBENS), Ecole Normale Supérieure, CNRS, INSERM, Université PSL, 75005 Paris, France; Institut de Biologie de l’ENS (IBENS), Ecole Normale Supérieure, CNRS, INSERM, Université PSL, Sorbonne University, Collège Doctoral, 75005 Paris, France; Université Paris Cité, CNRS UMR3738, Developmental and Stem Cell Biology Department, Institut Pasteur, Paris, France; CIRB, Collège de France, Université PSL, CNRS, INSERM, 75005 Paris, France

**Keywords:** in ovo electroporation, crispr/Cas9, knock-in, MMEJ, subcellular localization, endogenous tagging

## Abstract

Gene editing and protein tagging are at the heart of modern developmental and cell biology. The advent of CRISPR/Cas9 based methods offers the possibility to develop customized approaches for genomic manipulations in non-classical experimental models. Here, we show that highly efficient somatic knock-ins of long DNA fragments can be achieved in the developing chick neural tube in ovo. We compare different types of repair matrices and different methods for the delivery of the CRISPR/Cas9 machinery, and find that an all plasmid-based approach and short arms of homology provide an easy and efficient method to achieve high frequencies of knock-in insertions with virtually no background signal. We use this method to target fluorescent reporters and dynamically monitor the subcellular distribution of endogenously expressed tagged proteins, as well as to insert the Gal4-VP16 transcription factor or the Cre recombinase at specific loci to label neural sub-populations in the chick embryonic spinal cord. Finally, we show that the method can also be applied to target the epiblast and somitic mesoderm.

## Introduction

Avian models, and in particular the chick and quail embryos, have long been recognized as models of choice for developmental studies. One of the key advantages of these higher vertebrates is the direct accessibility to the embryo from very early developmental stages. Combined to the robustness of the embryo to manipulations, this has allowed seminal descriptive and functional studies throughout the 20st century. However, during the last decades, several limitations of avian models have hampered their large use: the advent of integration transgenesis, ES cell culture and methods for introduction of knock-out and knock-in modifications by homologous recombination have put the mouse model at the forefront of developmental biology (Aizawa, 2008), while gene manipulation in birds remained cumbersome; rearing large animals like chicken is not convenient and not accessible for most laboratories – although over the last decade, the comparatively smaller quail model has emerged as a convenient alternative model for the production and maintenance of transgenic animals bearing fluorescent reporters (Barzilai-Tutsch et al., 2022; Huss and Lansford, 2017; Serralbo et al., 2020); as a consequence, many more efforts have been invested in the development of tools, including specific antibodies, for studies in mammalian, and particularly in mouse models; in parallel, the emergence of zebrafish as a convenient, transparent model with strong genetic potential has led many groups to shift from the chick model to zebrafish for early developmental studies; however, birds occupy a strategic position in the phylogenetic tree and remain a key model for developmental and evo/devo comparative studies. In addition, they offer a number of interesting assets for experimental approaches. The accessibility and robustness of the embryo has always been an advantage for early embryogenesis, including but not limited to historic grafting experiments. In particular, viral vectors were developed that allowed somatic transfection and gene expression (Cepko et al., 1997) and were completed in the late 90s by the development of powerful in ovo electroporation methods that made “transient” somatic gene expression from simple plasmid transfection very easy in different structures of the embryo (Itasaki et al., 1999). Transposon-based integration of transgenes has recently made stable long-term expression possible (Loulier et al., 2014; Sato et al., 2007; Watanabe et al., 2007), and conditional systems like the Doxycycline-regulated Tet-On and Tet-off allow temporal regulation of transgene expression (Watanabe et al., 2007). With this arsenal of methods, the avian embryo remains a powerful model for functional studies in early development, at stages and in tissues that are not directly accessible in mammalian models. One of the main limitations remains the highly mosaic nature of somatic transgenesis: 1) only a subset of cells is targeted by electroporation methods (typically, less than 50% in the neural tube) and 2) the expression level of transgenes is highly variable due to differences in plasmid copy number being taken up by the cells. The latter point is particularly problematic for at least two approaches: in functional studies relying on dominant (gain or loss of function) approaches, the severity of the phenotype may differ depending on the number of copies of the transgene that are expressed; and when using expression of tagged constructs to monitor their subcellular distribution and dynamics, aberrant expression levels (compared to the endogenous protein) may result in localization artefacts, either due to aggregation or to saturation of binding partners necessary for subcellular targeting. Mosaicism in the number of active copies in a cell can be partially mitigated by the use of an elegant system based on conditional activation of expression of a transposon-based transgene (Kumamoto et al., 2020), however the expression levels still depend on insertion site and on the promoter used in these constructs. The other limitation is the low specificity of available tools for expression of transgenes in specific subsets of cells, although some studies have successfully used in ovo electroporation for the detailed study of cis-regulatory elements.

The advent of CRISPR/Cas9 as a highly efficient programmable gene edition tool holds the promise to facilitate genetic studies in virtually any species (Cong et al., 2013; Mali et al., 2013). Recent studies in the chick have already shown its potential to knock-out gene function, using the high frequency of small insertions/deletion induced by the Non-Homologous End Joining repair pathway (NHEJ) upon targeting by a gene-specific gRNA (Di Pietro et al., 2017; Gandhi et al., 2017; Véron et al., 2015). RNA-guided gene expression modulation has also been achieved in this model (Williams et al., 2018). More recently, knock-ins of reporters in genes of interest in the developing chick retina have been achieved via electroporation of Cas9/gRNA ribonuclear complexes and single strand DNA repair matrices. The specificity of reporter expression was apparent from the adequation between the expression of the reporters and the known pattern of expression of the targeted gene, demonstrating that it is a promising approach, although the efficiency has not been thoroughly quantified in these reports (Yamagata and Sanes, 2021; Yamagata et al., 2021). Interestingly, the authors reported that they obtained little success in the retina when they used plasmid-based expression of the Cas9 and gRNA components, compared to the use of in vitro preassembled Cas9 and guide RNA riboprotein complexes. Here, we explore the possibility of creating somatic knock-in using a combination of plasmid-based CRISPR/Cas9 targeting and repair constructs in the developing chick embryo. Primarily focusing on the early spinal cord neural tube, we show that highly efficient knock-ins can be achieved via simple in ovo electroporation. We compare the relative advantages of short and long arms of homology, and of circular or linear recombination matrices. In particular, we find that Microhomology-Mediated End Joining (MMEJ) based knock-in offers an optimal combination of efficiency and ease of use, since targeting reagents can be constructed without the need to obtain any genomic sequences. We show that the high efficiency of somatic knock in is compatible with multiplex gene tagging experiments. Targeting a number of genes, we show that tagging endogenous proteins via Crispr/Cas9-mediated knock-in recapitulates the subcellular distribution of the untagged protein more faithfully than overexpression of tagged proteins; tagging with fluorescent reporters opens the way to live monitoring of the subcellular dynamics of endogenous proteins with superior signal to noise ratio. We show that the Gal4-UAS and Cre-loxP binary expression systems can also be used for conditional expression of reporters in specific cell populations defined by the expression of the targeted gene, opening the way to gene-specific lineage tracing approaches. Due to the powerful signal amplification of conditional binary expression systems such as Gal4 or Cre, it has been suggested in previous reports in the mouse cortex that their use for somatic knock-ins might be prone to high unspecific background signal resulting from episomal leakiness (Tsunekawa et al., 2016). We show that the MMEJ approach efficiently suppresses the background signal resulting from unwanted episomal expression of the reporters. Finally, we demonstrate that somatic knock-in works, albeit with different efficiency, in other chick embryonic tissues. We anticipate that it will work with similar success in other avian species.

## Results

### Highly efficient introduction of a C-terminal GFP tag in the b-actin locus via HDR

We first tested a homology-directed repair (HDR) strategy (Figure 1A). We used a three-component system, based on simultaneous introduction of a homologous recombination vector (thereafter referred to as donor vector), a chimeric guide RNA and the spCas9 protein. The gRNA and spCas9 were carried by a single vector (Cong et al., 2013)(thereafter referred to as the Cas9-gRNA vector). We designed donor vectors using 1kb arms of homology flanking monomeric EGFP (mEGFP, thereafter GFP), NeonGreen (mNeonGreen, thereafter NG) or Cherry (mCherry, thereafter Ch) and tandem dTomato (tdTom, thereafter Tom) reporters, inserted as a fusion protein at the C-terminus of the ubiquitously expressed beta-actin gene (ACTB1, thereafter ACTB) sequence, one of the most highly expressed genes in transcriptomic data. Three guide RNAs targeting sequences near the stop codon of ACTB were chosen, using the CRISPOR design web site (http://crispor.gi.ucsc.edu/). As negative control, a sequence that does not target any region in the chick genome (Gandhi et al., 2017) was chosen (see Methods).

**Figure 1.**
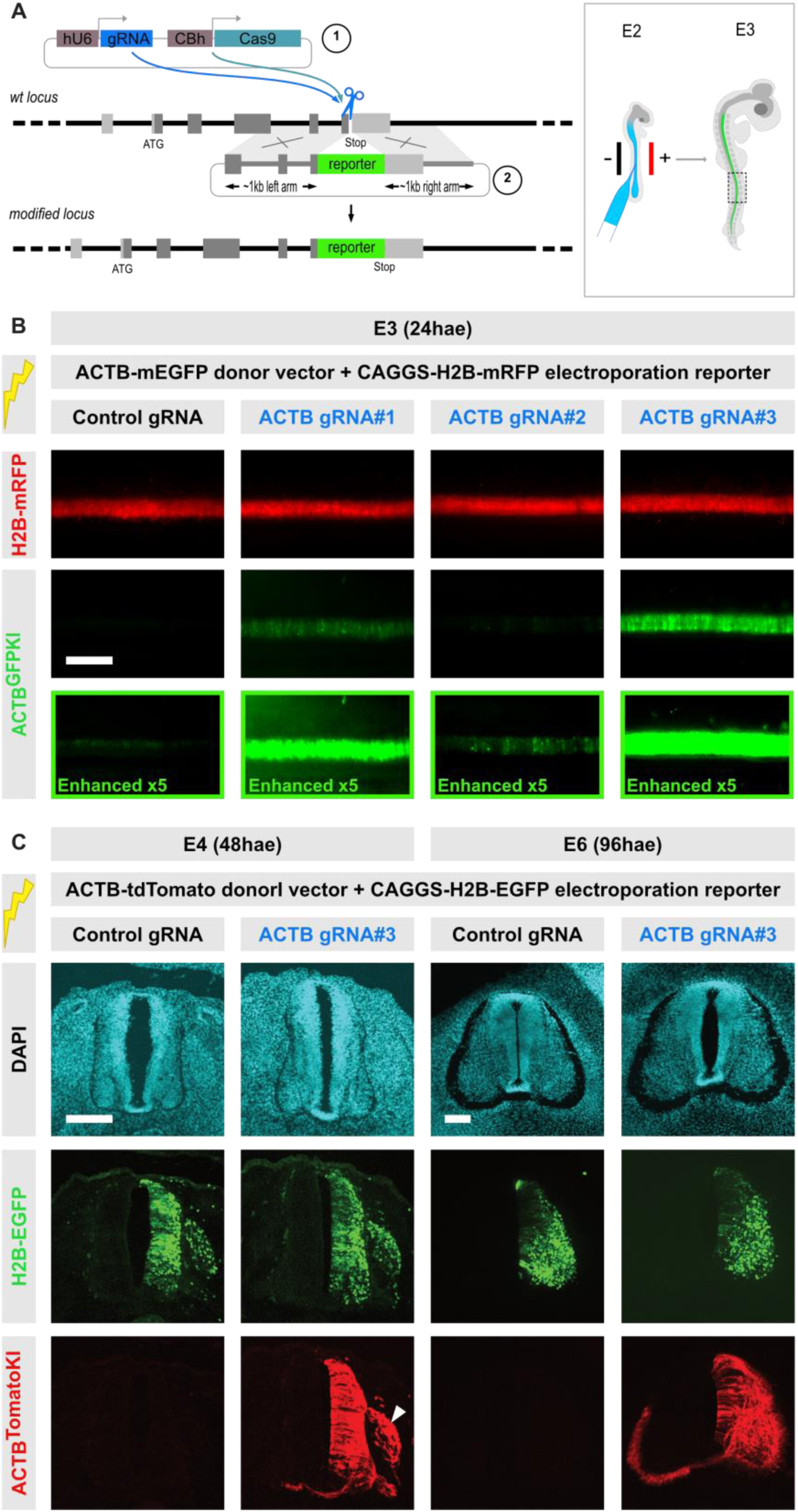
– An all-plasmid based strategy for somatic knock-ins in the chick embryonic neural tube. A. Principle of the HDR knock-in strategy. The locus of interest is targeted by a Cas9-gRNA vector (1) expressing a locus-specific gRNA (in the schematized example, which reproduces the chick ACTB locus organization, the gRNA targets the C-terminus of the coding sequence). The donor vector (2) contains ∼1kb long arms of homology chosen on both sides of the gRNA target region, and flanking a reporter gene sequence. Linearization of the target locus by vector 1 induces a DNA repair mechanism that uses homologous recombination with vector 2 to introduce the reporter sequence as a fusion to the coding sequence of the target gene (a C-terminal fusion protein in the example schematized here). B. Characterization of the knock-in efficiency at the ACTB locus using the strategy described in A. The ACTB-mEGFP donor vector was coelectroporated with an electroporation reporter (CX-H2B-mRFP) and a Cas9-gRNA vector expressing the Cas9 protein and either a negative control gRNA or one of three gRNAS targeting the C-terminal region of ACTB. Electroporation was performed at E2 (HH stage 14) and embryos showing similar electroporation levels (red) were imaged directly in ovo 24 hours later. Two of the three ACTB gRNAs show a high GFP signal, whereas the third one shows much fewer GFP-positive cells (only visible on the enhanced image in the bottom row). The negative control has virtually no GFP signal, showing that the signal observed in the three other conditions is specific of insertions at the target locus. Scale bar, 300µm. C. The ACTB-tdTomato donor vector was coelectroporated with an electroporation reporter (CX-H2B-EGFP) and a Cas9-gRNA vector expressing the Cas9 protein and either a negative control gRNA or gRNA#3 targeting the C-terminal region of ACTB. Transverse sections at the thoracic level of embryos harvested at E4 and E6, respectively 48 and 96 hours after electroporation at E2, show a very strong ACTB-Tomato signal along the whole DV extant of the neural tube of embryos electroporated with ACTB gRNA#3, and no signal in the control condition, despite similar electroporation level with the H2B-EGFP reporter. In addition to the neural tube, a strong knock-in signal is visible at E4 in the neural crest-derived dorsal root ganglia (white arrowhead). Scale bars, 150µm

The ACTB-GFP donor and Cas9-gRNA vectors were introduced in the chick embryonic neural tube at stage HH14 (52-54h of incubation, Embryonic day 2, E2) by in ovo electroporation (Figure 1A), together with a plasmid reporting electroporation efficiency through expression of a H2B-mRFP1 (thereafter RFP) fluorescent reporter driven by the ubiquitous CAGGS promoter (Niwa et al., 1991), thereafter referred to as CX in text and figures). Embryos were monitored for GFP signal 24 hours after electroporation. Whereas no signal could be detected in embryos that had received the negative control gRNA, a strong GFP signal overlapping with the RFP reporter was detected in the neural tube of embryos that had received ACTB gRNA (Figure 1B). While all three ACTB gRNAs yielded some GFP positive cells, one of them only produced a modest signal, while the other two resulted in strong GFP expression. This agrees with gRNAs harboring different targeting efficiencies. We obtained similar results with the ACTB-tdTomato matrix and an GFP electroporation reporter. In transverse sections (from gRNA#3) 48 hours (E4) and 96 hours (E6) after electroporation, the Tomato signal was detected in a large number of GFP positive cells (Figure 1C). Altogether, these experiments indicate that a donor plasmid designed to target a fluorescent protein fusion reporter to the C-terminus of the chick b-actin gene with 1kb long arms of homology is highly efficiently and specifically introduced by somatic homologous recombination to the endogenous locus in presence of a Cas9 and a locus-specific gRNA.

### Using short arms of homology (the Pitch method)

Having established that HDR-based somatic knock-in works in the neural tube, we proceeded to generate donor vectors based on the same design to target other genes of interest. The approach was successful in a number of genes (see below), but we were faced with several difficulties. For the construction of donor vectors, the PCR amplification of long arms of homology from GC-rich genomic loci can be difficult, even using optimized protocols. The same GC-rich sequences are also difficult to generate by gene synthesis techniques, and require expensive optimization. In addition, we anticipated that using ∼1kb arms of homology for insertion of a reporter at the N-terminus of a coding sequence would often involve the inclusion of promoter sequences in the construct, and in some cases lead to strong background expression of the reporter from the plasmid.

We therefore sought to evaluate alternative strategies. Single stranded DNA matrices with short arms of homology (<70 bases) work very efficiently for knock-in approaches in a number of cell types. This approach typically requires to simultaneously introduce Cas9 protein and the gRNA in the form of an in vitro preassembled ribonucleoprotein complex rather than in a plasmid encoded form: single stranded DNA matrices have a short half-life in the cell and will likely be destroyed before plasmid-based Cas9 and gRNA expression reaches sufficient levels in the cell to linearize the target locus. Indeed, Yamagata and Sanes have reported somatic knock-in using electroporation of single stranded DNA repair matrices in the chick embryonic retina when using a purified ribonucleic Cas9/gRNA complex, but virtually no knock-in events when Cas9 and gRNA were expressed from a plasmid (Yamagata et al., 2021). Single stranded DNA fragments up to 200bp can be readily synthetized by manufacturing companies, but this limits the size of the reporter insert to roughly 60 bases, restricting the approach to the use of classical short epitope tags (flag, HA, Myc) or to the “tag” moiety of split fluorescent proteins (Cabantous et al., 2005; O’Hagan et al., 2021; Zhou et al., 2020). Alternatively, longer single stranded fragments can be generated by asymmetric PCR, allowing the production of repair templates with longer insertions (typically, a full length GFP) (Yamagata et al., 2021). In our hands, the asymmetric PCR production of long ssDNA matrices was impractical, requiring time consuming and costly optimization steps and ultimately yielding low amounts of matrix from a lot of PCR material (not shown). We therefore tested the single stranded oligonucleotide approach and tested the efficiency of insertion of a ssDNA oligonucleotide carrying the GFP_11_ tag sequence from split-GFP (Cabantous et al., 2005) in the ACTB locus. To compare this method with our plasmid-only approach, we constructed a donor vector in which the GFP_11_ tag sequence is flanked with the same 1kb long arms of homology as above. To complement the GFP_11_ tag and produce a fluorescent signal, a vector expressing the GFP_1-10_ sequence under the strong CX promoter was constructed and included in the electroporation mix. Both approaches yielded specific knock-in signal (Supp Figure 1B and 1D), but the electroporation efficiency was consistently much higher in the plasmid-only approach compared to the ribonucleoprotein mix protocol, such that more knock-in events were recovered. As reported by Yamagata and Sanes, we also observed that combining the ssODN with plasmid-encoded gRNA and Cas9 was less efficient than a mix of ssODN and ribonucleic complex (Supp Figure 1C).

It has also been reported that 5’-end biotinylated double stranded PCR fragments can be used as repair templates in association with Cas9/gRNA ribonucleic complexes in different models (Paix et al., 2017; Paix et al., 2023; Seleit et al., 2021). We therefore generated a double-stranded GFP repair matrix targeting the ACTB locus based on this design. However, we found that this approach also gave poor results compared to the all plasmid approach with the ACTB-GFP donor (Supp figure 2).

The low efficiency of the approaches based on preassembled ribonucleic complexes presumably derives mainly from the low efficiency of co electroporation of positively charged Cas9 protein with negatively charged nucleic acids, rather than from the ability of linear DNA fragments with short homology to be used as repair templates. On the other hand, the even lower efficiency of the combination of linear single or double stranded donor fragments when combined with plasmid-based expression of gRNA and Cas9 probably results from the rapid degradation of the linear fragments before production of the Cas9/gRNA complexes in electroporated cells. We therefore tested whether combining the production of a linear repair template inside the cell from an electroporated circular plasmid combined with the efficient introduction of plasmid-based Cas9 and gRNA would circumvent these limitations. We adapted the CRIS-PITCh (CRISPR based Precise Integration into Target Chromosome) strategy that has been used successfully in a number of model systems and cell lines (Aida et al., 2016; Almeida et al., 2021; Ezaki et al., 2022; Hisano et al., 2015; Nakade et al., 2014; Sakuma et al., 2016; Welker et al., 2021; Wierson et al., 2020). Briefly, the PITCh approach relies on Microhomology-Mediated End Joining (MMEJ) rather than HDR to mediate repair, and uses short arms of homology (usually 20-35 bases) from a linearized template. The template is excised in cells from a circular donor vector, typically through CRISPR/Cas9 mediated linearization via generic gRNA target sites that flank the arms of homology (Nakade et al., 2014).

We constructed a donor vector containing a NeonGreen reporter flanked with 35bp homology sequences targeting the ACTB C-terminal region, flanked by two “universal” gRNA target sites (Figure 2A). The universal gRNA (thereafter referred to as uni2 gRNA, or uni2) has previously been described in a study in zebrafish (Welker et al., 2021), which shows its high efficiency in vector linearization, and it does not target any sequence in the chick genome. We then evaluated the MMEJ approach and compared its efficiency with the HDR approach described above. As shown in figure 2B, both MMEJ and HDR yielded a strong and specific signal when targeting the ACTB locus with the same gRNA targeting the ACTB C-terminus, and absolutely no signal with a negative control gRNA. In contrast to HDR, in which the long arms of homology of the circular vector are sufficient for homologous recombination at the linearized genomic locus, the MMEJ approach absolutely required release of the linear fragment from the circular vector, and no signal was observed when the linearizing gRNA was absent (Figure 2B and Supp Figure 3C-C’).

**Figure 2.**
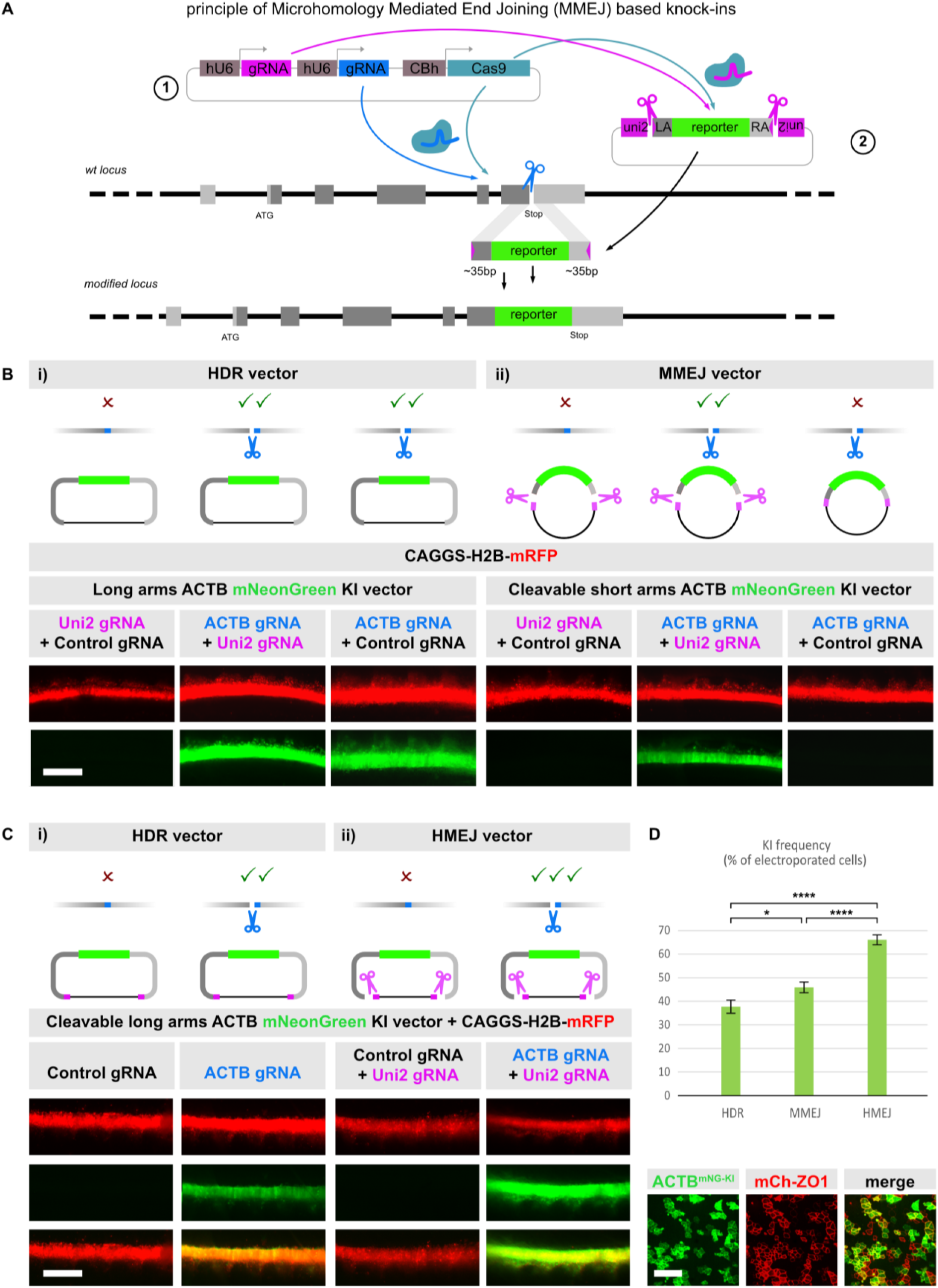
– CRISPR/Cas9-based post-transfection linearization of the donor vector improves knock-in efficiency. A. Principle of microhomology-mediated end joining (MMEJ) based knock ins. The locus of interest is targeted by a Cas9-gRNA vector (1) expressing a locus-specific gRNA (in the schematized example, which reproduces the chick ACTB locus organization, the gRNA targets the C-terminus of the coding sequence). In this scheme, vector 1 also expresses a highly efficient “universal” (uni2) gRNA that does not target any sequence in the chick genome. The donor vector (2) contains short ∼35bp arms of homology chosen on both sides of the gRNA target region, and flanking a reporter gene sequence. The arms of homology are flanked by two uni2 gRNA target sequences that allow linearization of vector 2 by the uni2gRNA/Cas9 complex encoded by vector 1, and release of a linear double stranded donor template. In electroporated cells, simultaneous linearization of vector 2 and the target locus induces a DNA repair mechanism that uses the microhomology between the exposed ends of the released donor fragment and the genomic DNA to introduce the reporter sequence as a fusion to the coding sequence of the target gene (a C-terminal fusion protein in the example schematized here). B. Comparison of HDR (i) and MMEJ (ii) strategies for knock-in of a mNeonGreen reporter at the C-terminus of the ACTB locus. For the MMEJ strategy (ii), a signal is only observed when the two gRNAs that target the genomic locus and the donor vector are provided (ACTB + uni2 gRNA, center panel). No signal is obtained when only the donor vector (uni2 + control gRNA, left) or only the genomic locus (ACTB + control gRNA, right) are targeted. Scale bar, 300µm. C. Comparison of HDR (i) and HMEJ (ii) strategies for knock-in of a mNeonGreen reporter at the C-terminus of the ACTB locus. Note that when the ACTB gRNA is present and KI events are observed, the relative intensity of the green compared to the red channel appears stronger with the linearized donor vector (HMEJ, ACTB + uni2 gRNA condition, right column in ii) compared to the circular donor vector (HDR, ACTB gRNA alone, right column in i). Scale bar, 300µm. D. Quantification of the knock-in efficiency of a mNeonGreen reporter at the C-terminus of the ACTB locus with the three different knock-in strategies. Embryos were electroporated at E2 and harvested at E3. A plasmid expressing a mCherry-ZO1 reporter was used to evaluate electroporation quality, and KI efficiency was measured as the ratio of mNeonGreen-positive to mCherry-ZO1 positive cells in en-face views or the apical surface of open-book mounted neural tubes. A representative image from the HMEJ condition is shown at the bottom of the panel. Quantifications are from 6 fields of view from 4-6 embryos for each condition, for a total of >2900 electroporated cells/condition. Unpaired Students t-test, *: p=0.0333; ****: p=0.0001. Scale bar, 10µm.

As controls for specificity, we generated constructs bearing only one linearization site, or constructs bearing the two linearization sites but either only one (left or right) arm, or no arm of homology. Supplementary Figure 3E-I shows that having only one arm of homology exposed can result in rare GFP positive cells, suggesting that asymmetric repair may occur at very low frequency via a combination of MMEJ on one arm and Non-Homologous End Joining (NHEJ) repair on the other arm. On the other hand, linearized vectors releasing a reporter devoid of arms of homology yielded absolutely no signal (Supplementary Figure 3D), demonstrating the inefficiency of a NHEJ-based strategy for knock-in insertions in the neural tube. Overall, this indicates that MMEJ is efficient and highly specific, and that it requires two exposed arms of homology with the two regions flanking the genomic target site.

### Long arms PITCH – HMEJ

In a recent paper evaluating different lipofection methods to transfect chick primordial germ cells (PGCs) in vitro, Watanabe and colleagues compared the frequency of knock-in events of a GFP reporter at the ACTB C-terminus using a “classical” circular HDR vector that uses 500bp arms of homology, and a modified version of this vector that can be linearized thanks to the addition of gRNA target sites flanking the arms of homology (Watanabe et al., 2023). This approach that combines the long arms of homology of HDR with the linearization used for MMEJ is referred to as HMEJ (Homology-Mediated End Joining). Although knock-in frequencies are extremely low in PGCs, this study reported a 3 to 10-fold increase in recombination frequency with the linear (HMEJ) compared to circular (HDR) vector. We wondered whether HMEJ would improve knock-in frequency over HDR and MMEJ in the chick spinal cord. To test this possibility, we flanked the 1kb arms of homology in our ACTB-NeonGreen HDR vector with uni2 target sites. We also generated dual gRNA vectors (Cas9-dgRNA vectors) that bear two gRNA cassettes driving the uni2 gRNA and either the ACTB or a control gRNA under human U6 promoter, and Cas9 under the CBh promoter (as depicted in Figure 2A). The HDR and HMEJ vectors were electroporated with either control or ACTB dgRNA vectors, and a CX-H2B-RFP electroporation reporter. 24hours after electroporation, both strategies yielded a strong and specific NeonGreen signal, but the NeonGreen signal consistently appeared stronger (Figure 2C) in the HMEJ compared to HDR condition, suggesting a higher frequency of knock-ins with HMEJ. To directly compare the efficiency of the three approaches, the HDR, MMEJ and HMEJ vectors were each electroporated together with the ACTB dgRNA vector, and a CX-Cherry-ZO1 electroporation reporter. 24 hours after electroporation, embryos were harvested, fixed, the neuroepithelium was dissected and mounted in an open-book configuration for en-face imaging of the apical surface. To evaluate the knock-in efficiency of the three approaches, we evaluated the percentage of ACTB-NeonGreen positive cells compared to Cherry-ZO1 cells in confocal images at the level of apical junctions of neural progenitors. We used automated segmentation with Cellpose2 (Pachitariu and Stringer, 2022), with minimal visual supervision and manual correction. HDR, MMEJ and HMEJ yielded on average 37.8+/−2.9 %, 45.9 +/− 2.3 %, and 66.3+/− 2.1% of knock-in events, respectively (Figure 2D), confirming that combining long arms of homology and donor vector linearization in the HMEJ approach improves over the two parameters used separately in HDR and MMEJ. We did not evaluate intermediate arm length between the 35bp of the MMEJ construct and the 1kb of the HMEJ. However, in early tests with circular HDR vectors at the ACTB and other loci, we obtained similar efficacy with 500bp and 1kb arms (not shown), suggesting that the optimal arm length for a HMEJ approach would be lower than 500bp.

### Efficient biallelic and dual gene targeting with HMEJ

The high frequency of knock-in achieved with the three approaches, and most impressively with HMEJ, suggests that it should be possible to obtain bi allelic events or to target two –or more-genes simultaneously at high frequency with these approaches. To evaluate the frequency of biallelic knock-ins in HDR and HMEJ, we constructed an ACTB-Cherry HMEJ donor vector. We electroporated embryos with equal amounts of ACTB-NG and ACTB-Cherry donors, in combination with either the single Cas9-gRNA vector targeting only ACTB (mimicking an HDR condition) or the dual Cas9-gRNA vector targeting ACTB and the uni2 sequence, and an electroporation reporter (CX-iRFP-ZO1). As expected, linearization of the vectors (HMEJ) yielded a higher frequency of knock-in events (70.1 +/−1.4 %) than using circular plasmids (HDR, 47.5+/−2. 6 %), measured as the total percentage of “green” and/or “red” cells in the iRFP-ZO1+ electroporated population. Importantly, the percentage of double positive cells was visibly higher with HMEJ (Figure 3A) and rose from 4.9+/−0.6% with circular vectors to 17+/−2.1% with linear vectors (Figure 3B). As double color (red AND green) biallelic insertions are as likely to occur as single color (red OR green) biallelic insertions, this suggests that the frequency of biallelic insertions rose from 9.8% with circular vectors to an impressive 34% of electroporated cells when using linearized vectors (Figure 3C). Importantly, the background was null in the absence of the ACTB gRNA: using a dual Cas9/gRNA vector that expresses the negative control and uni2 gRNAs, we observed only one cluster of 4 GFP-positive cells, and one pair of Cherry-positive cells out of several thousand iRFP-ZO1-positive cells from 4 embryos. The subcellular distribution of the GFP and Cherry positive signal in these rare cell clusters was the same as in the knock-in conditions, with an enrichment near apical junctions, suggesting that these cells may correspond to spontaneous homologous recombination events at the ACTB locus (not shown).

**Figure 3.**
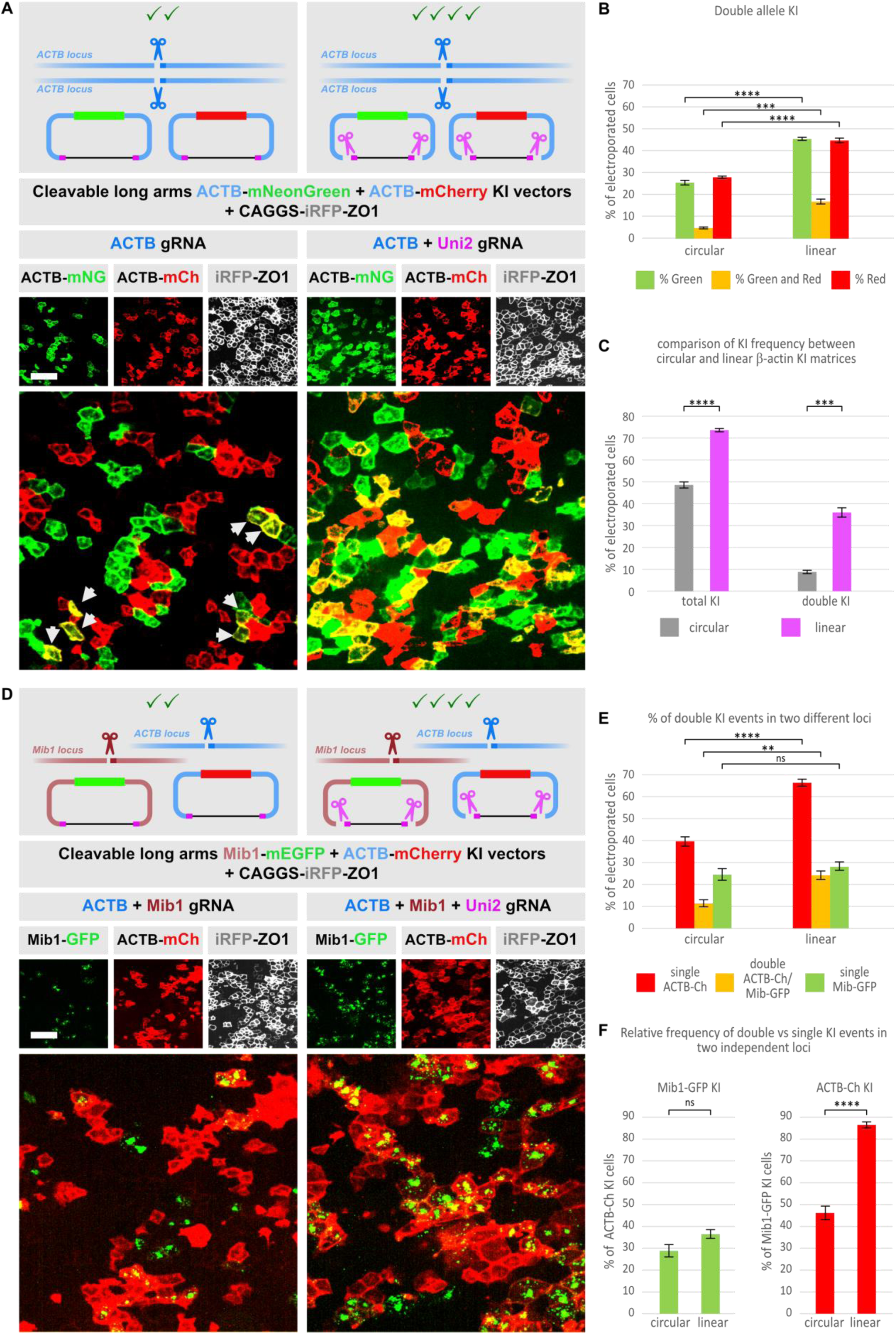
– Efficient double-allele and double-gene targeting with linearized donor vectors. A. Comparison of the efficiency of circular HDR (left) and linear HMEJ (right) vectors for double allele knock-in insertion. The ACTB locus was targeted with a mix of vectors introducing mCherry and mNeonGreen fluorescent reporters at the C-terminus of ACTB coding sequence. A gRNA targeting ACTB was introduced alone (left) or in combination with the uni2 gRNA that linearizes the targeting vector (right). A iRFP-ZO1 expression vector was used as electroporation reporter. Representative en-face views of the neuroepithelium 24 hours after electroporation are presented. Only few double positive cells (yellow cells, pointed with white arrows) are observed in the HDR condition. Vector linearization increases the overall frequency of KI events, leading to a massive increase in the frequency of simultaneous insertion of the mCherry and mNeonGreen cassettes at two alleles (yellow cells). Scale bar, 10µm. B. Graphic representation of the percentage of iRFP-ZO1-electroporated cells with single and dual color ACTB knock-in insertions upon targeting with circular and linear vectors. Unpaired Student’s t-test, ***: p=0.0013; ****: p<0.001. C. Graphic representation of the percentage of iRFP-ZO1-electroporated cells with ACTB knock-in insertions (light blue) and deduced percentage of double allele insertions (dark blue) upon targeting with circular and linear vectors. The percentage of double insertions is twice the percentage of “green and red” cells in panel B. Unpaired Student’s t-test, ***: p=0.0013; ****: p<0.001. D. Comparison of the efficiency of circular HDR (left) and linear HMEJ (right) vectors for simultaneous knock-in insertion in two independent loci. The Mib1 and ACTB loci were targeted with a mix of vectors introducing mCherry at the C-terminus of ACTB and mEGFP at the C-terminus of Mib1 coding sequences. gRNAs targeting ACTB and Mib1 were introduced without (left) or in combination with the uni2 gRNA that linearizes the targeting vectors (right). A iRFP-ZO1 expression vector was used as electroporation reporter. Representative en-face views of the neuroepithelium 24 hours after electroporation are presented. Vector linearization increases the overall frequency of KI events, leading to a massive increase in the frequency of simultaneous insertion of the mCherry and mEGFP cassettes at the two genes (yellow cells). Scale bar, 10µm. E. Graphic representation of the percentage of iRFP-ZO1-electroporated cells with a single (green or red) or both (yellow) genes successfully targeted with circular and linear vectors shows a massive increase in the frequency of double gene targeting upon vector linearization. Unpaired Student’s t-test, **: p=0.0023; ****: p<0.001. F. Graphic representation of the percentage of cells showing a Mib1 KI event within the population of cells with ACTB knock-in insertions (green, left) and of the percentage of cells showing an ACTB KI event within the population of cells with Mib1 knock-in insertions (red, right). Unpaired Student’s t-test, ****: p<0.001.

To test the ability of the method to achieve knock-ins in two different genes simultaneously, we constructed a vector targeting the Notch pathway regulator Mindbomb1 (Mib1). We have previously shown that Mib1 is expressed in neural progenitors at embryonic day 3, and that it is located at pericentriolar satellites during interphase (Tozer et al., 2017). Knock-in of a fluorescent reporter was therefore expected to produce a signal near the apical surface of neuroepithelial progenitors, such that knock-in efficiency would be easy to monitor and quantify in en-face views of the neuroepithelium. We generated versions that tag the C-terminus of Mib1 with GFP, using 1kb arms of homology flanked by uni2 target sites. We co-electroporated the Mib1-GFP and ACTB-Cherry repair vectors together with vectors expressing Cas9 protein and gRNAs targeting Mib1, ACTB, and with or without the uni2 gRNA to test single and double knock-in efficiencies under HMEJ or HDR conditions. Again, the CX-iRFP-ZO1 vector was added as an electroporation reporter. Embryos were electroporated at HH stage 14 and harvested 24 hours later, and the neuroepithelium was dissected for en-face imaging of the apical surface.

As for dual allele knock-in above, a striking increase in the frequency of dual gene insertions with linear compared to circular vectors was immediately obvious in en-face images (Figure 3D). In the HDR (circular vector) condition, we counted 39.7+/−3.9% of ACTB-Cherry+ cells and 24.8+/−4.9% of Mib1-GFP+ cells, with 11.4+/−2.9% of cells having insertions in both genes (Figure 3E). Under HMEJ (linearized vector) conditions, the percentage of ACTB-Cherry+ cells rose to 66.5+/−3%, while the increase in the number of Mib1-GFP+ cells was not significant (28+/−3.5%). Strikingly, 24.3+/−3.5% of the electroporated cells had insertions in both genes (Figure 3E). In particular, 86.5+/−2.6% of the Mib1-GFP+ cells also displayed an ACTB-Cherry insertion in the linearized condition (Figure 3F).

Overall, these experiments show that targeting two independent genes can be achieved at high frequency with HMEJ vectors. We did not evaluate the efficiency of the MMEJ approach for dual allele or dual gene targeting. We anticipate that it might be slightly lower than with HMEJ, as MMEJ was slightly less efficient than HMEJ in our single-color experiments targeting the ACTB locus, but that it would still yield high numbers of dual targeting. Importantly, the efficiency of dual gene targeting will depend on the efficiency of targeting for each gene: in addition to the chosen targeting method (HDR, MMEJ and HMEJ), this is expected to vary between genes and will also depend on the identification of efficient gRNAs for each locus.

### Endogenous protein tagging for dynamic live reporting of protein subcellular distribution

Our initial drive for using somatic knock-in was to monitor the subcellular localization of proteins of interest. Subcellular localization of proteins can be documented via specific antibodies directed against epitopes in the protein. While this gives access to the endogenous protein, it is highly dependent on the availability of specific antibodies, which can be tricky to develop. Most commercial antibodies are directed against proteins from the most frequently used human and mouse models, and cross-reactivity to distant models such as birds is usually a matter of chance; besides, fixation of the samples is necessary, and fixation conditions can affect the final results. Alternatively, exogenous expression of tagged proteins overcomes the need for specific antibodies, and fluorescent tags allow monitoring in live conditions. However, it is difficult to control the expression levels of tagged proteins. Besides, exogenous tagged proteins compete with the endogenous untagged version: in cases where subcellular distribution relies on limiting availability of binding partners, this can result in localization artefacts, or lead to abnormal aggregation when exogenous proteins largely exceed endogenous concentrations. In this context, tagging proteins expressed at physiological levels from their endogenous locus vi a knock-in approach appears as a superior alternative, and the method of choice for the documentation of dynamic cellular processes in developmental contexts. In the following section, we illustrate the efficient tagging of several endogenous proteins. We present examples using either circular HDR or linear MMEJ donor vectors, as these data on protein tagging were obtained in parallel to our exploration of the efficiency of the different methods.

We started by targeting the Mindbomb1 (Mib1) locus for C-terminal GFP fusion. We have previously described that Mib1 localizes to centriolar satellites in neuroepithelial cells, using both an anti-Mib1 antibody and moderate overexpression of tagged Mib1 (Tozer et al., 2017). The Mib1 antibody was very sensitive to fixation conditions, leading to reproducibility issues and often yielding high background; on the other hand, overexpression resulted in variable expression levels and, while Mib1 localization at satellites appeared to withstand robust overexpression without saturation at satellites, we frequently observed aggregates of the tagged protein in cells with strong expression, which may be attributed to an overexpression-driven enhancement of Mib1’s natural ability to multimerize (Cao et al., 2024). Besides, since satellites and centrosomes are liquid-liquid phase separated organelles whose size and organization relies on the levels of their core components, there is a significant risk that overexpression of some of their components might imbalance their organization. Therefore, we were pleased to observe that endogenous Mib1-GFP recapitulated the localization at centriolar satellites that we previously described (Fig 4A). Mib1-GFP signal intensity at apical satellites was much more homogeneous with the knock-in strategy than with overexpression and also produced less aggregate-like signal in the tissue (not shown). Using live imaging, we found that the Mib1-GFP knock-in recapitulated the specific mitotic asymmetry at spindle poles that we previously described in dividing neuroepithelial progenitors (Figure 4B). We also tested several other tags in fusion to endogenous Mib1, including mRuby3, tdTomato, SNAP-Tag and the recently developed pFAST (Benaissa et al., 2021), and observed the same localization at satellites with each of them. pFAST is a ligand-dependent fluorogenic tag that is shorter than most fluorescent tags and whose emission wavelength can be tuned from Cyan to Red through the use of different cell-permeant ligands. Using HMBR (green) or HBR3.5DM (red) ligands, we show here that a single knock-in matrix can produce signals at different wavelengths. In particular, Mib1-pFAST with the HMBR ligand produced a signal of similar intensity as Mib1-GFP (Figure 4A).

**Figure 4.**
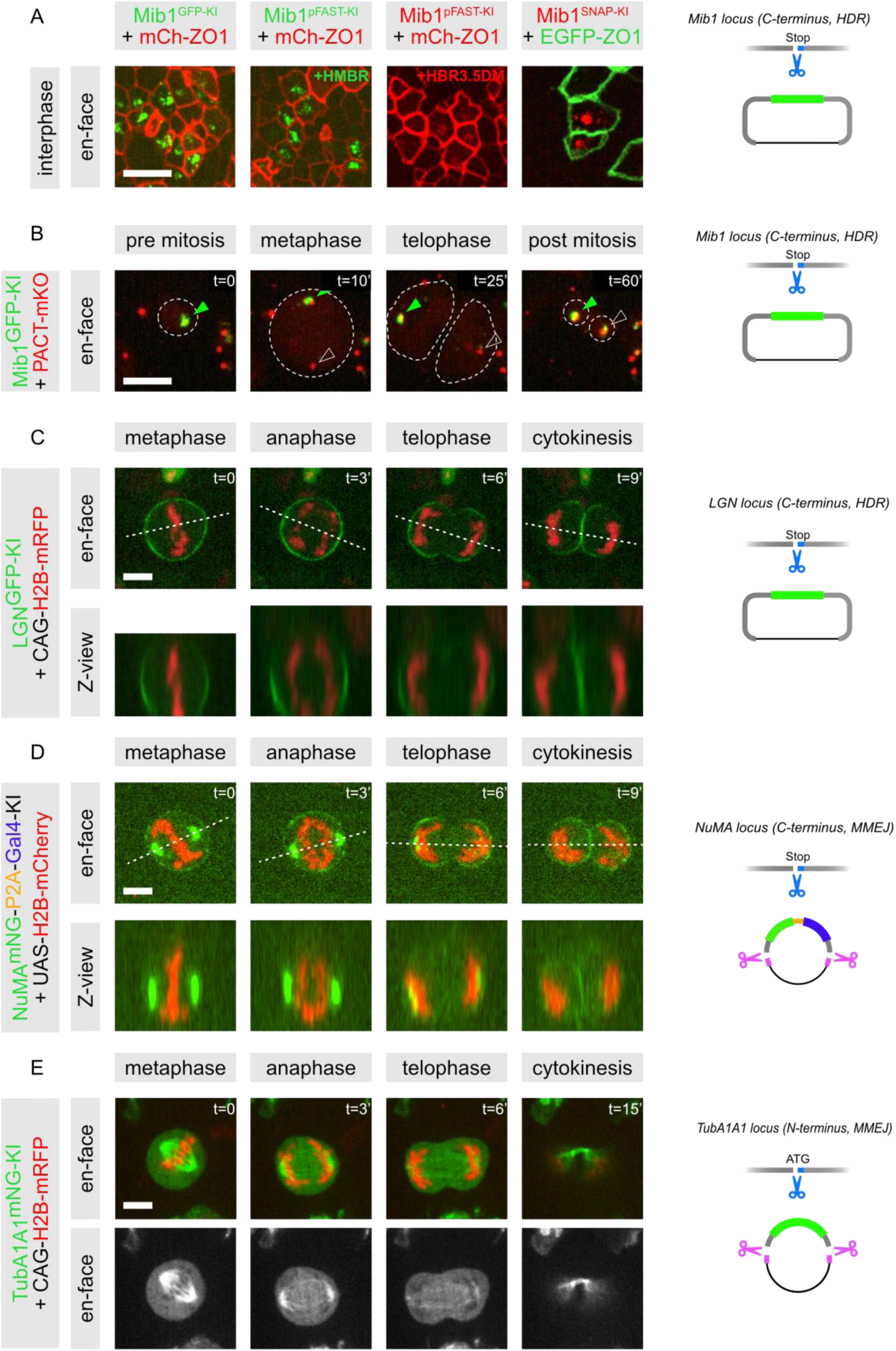
– Live monitoring of the subcellular distribution of endogenously tagged proteins via somatic knock-in of fluorescent reporter. The subcellular localization of fusion fluorescent reporters faithfully reproduces the expected distribution of the proteins expressed from targeted loci, and can be monitored in live imaging conditions at endogenous expression levels. Panels B, C, D, E show time lapse sequences of the distribution of tagged Mib1, LGN, NuMA and Tubulin alpha1A in dividing cells. Panel A illustrates the apical distribution in interphase neuroepithelial cells of Mib1 tagged with different reporters (EGFP, pFAST, SNAP-Tag). The apical junctional complexes are visualized via coelectroporation of mCherry-ZO1 or mEGFP-ZO1 reporters. Panel B shows the asymmetric distribution of Mib1-GFP on one pole of the mitotic spindle during mitosis at E3, confirming previous reports with an overexpressed Mib1-GFP (Tozer et al, 2017). Spindle poles are visualized via the co-elecroporation of a PACT-mKO1 reporter (red). Panel C shows the cortical distribution of LGN-GFP during mitosis of neuroepitheial progenitors at E3. Z-views show a lateral enrichment at the cell’s equator from metaphase to telophase, confirming previous reports from fixed material (Saadaoui et al, 2014), and indicates a relocalization at the interface between sister cells after cytokinesis. Progression through phases of mitosis is visible via the co-electroporated vector expressing the H2B-mRFP reporter under control of a ubiquitous reporter (red). Panel D shows the cortical distribution of NuMA-mNeonGreen during mitosis of neuroepitheial progenitors at E3. Z-views show a lateral enrichment at the cell’s equator from metaphase to telophase, confirming previous reports from fixed material (Peyre et al, 2011). The knock-in constructs contains a P2A-Gal4 cassette immediately downstream of the mNeonGreen fluorescent reporter, allowing the production of the Gal4 transcription factor in KI-positive cells and specific activation of a UAS-H2B-mCherry transgene in these cells. Panel E illustrates the insertion of a mNeonGreen fluorescent reporter at the N-terminus of the TUBA1A1 locus, and shows the specific staining of the microtubule network during mitosis of a neuroepithelial cell. Note the bright contrast of the mNeonGreen TUBA1A1 signal between the mitotic spindle and the cytoplasm. The right part of the figure illustrates the diverse architectures of the donor vectors: circular HDR versus linear MMEJ, C-terminal versus N-terminal targeting, and addition of the binary Gal4 system for activation of an independent reporter in knock-in cells. Scale bars, 5µm

We went on to tag the G-protein regulator LGN (GPSM2), a key regulator of mitotic spindle orientation in multicellular organisms (reviewed in (Di Pietro et al., 2016). In epithelial structures, LGN localizes to the lateral cortex of mitotic cells where it recruits dynein motors via the Nuclear and Mitotic Apparatus (NuMA) adapter protein, and generates localized forces that “pull” on astral microtubules to orient the mitotic spindle parallel to the plane of the epithelial surface (Di Pietro et al., 2016). While the mechanisms that restrict LGN to the lateral cortex are not completely understood, interaction of LGN’s C-terminal GPR domains with cortical Gai subunits in their GDP-bound conformation is essential. The availability of GDP-Gai appears as a limiting factor at the cortex, and endogenous LGN levels in the cell are low. As a result, we have previously observed that expression of GFP-tagged LGN saturates its cortical interactors, leading to artificial accumulation of cytoplasmic GFP-LGN that shadows the specific cortical localization, even at moderate expression levels, making it impossible to document live LGN subcellular dynamics in the neuroepithelium (Saadaoui et al., 2014; Saadaoui et al., 2017). We generated a GFP knock-in matrix targeting the C-terminus of LGN, and coelectroporated this matrix with relevant Cas9-gRNA vectors and a H2B-mRFP reporter. In parallel, we electroporated our previously described CMV driven GFP-LGN overexpression vector together with H2B-mRFP (Saadaoui et al., 2014). Embryos were harvested and fixed 24hae, processed for whole neural tube staining with anti-GFP and anti-LGN antibodies, and mounted for en-face imaging. In the knock-in embryos, we observed numerous mitotic cells with highly contrasted cortical GFP staining, that mirrored the anti-LGN staining. Overall, cortical GFP intensity was homogenous between cells, as expected for signal resulting from knock-in insertions at the LGN locus. In contrast, in cells expressing the CMV-driven GFP-LGN fusion, GFP signal intensity in mitotic cells was highly variable, consistent with variable plasmid copy number between cells. Cells with exogenous GFP-LGN fluorescence levels comparable to endogenous (knock-in) LGN-GFP generally displayed a lower cortical GFP intensity than knock-in cells, whereas cells with higher fluorescence levels clearly had strong cytoplasmic LGN accumulation, as evidenced by both GFP and LGN antibody stainings (Supplementary Figure 4). Taking advantage of GFP fluorescence and of the improved contrast compared to overexpression, we were able for the first time to monitor LGN dynamics in dividing neuroepithelial progenitors through en-face live imaging on a spinning disk confocal microscope (Figure 4C). The mosaic nature of the knock-in provided the additional advantage that isolated positive cells could be monitored, allowing to precisely assign the cortical signal to the positive cell rather than to its neighbors, which can be tricky with antibody staining. This allowed to visualize the lateral enrichment of LGN in lateral views of dividing progenitors (Figure 4C, bottom panels). Notably, we observed that upon cytokinesis, LGN is rapidly relocated from its lateral cortical distribution to the cortical interface between the two newborn sisters.

We used the MMEJ strategy to insert a NeonGreen tag at the C-terminus of the LGN partner NuMA. NuMA is normally nuclear during interphase, and relocates to the mitotic apparatus during mitosis, but also interacts with LGN at the lateral cellular cortex. We introduced a P2A pseudo-cleavage sequence followed by the Gal4-VP16 (thereafter Gal4) transcription factor downstream of the NeonGreen fusion tag. Adding Gal4 permits to replace the ubiquitous electroporation reporter that was used in our previous examples with a UAS-driven reporter that will only mark cells with a knock-in event, provided that they express the targeted gene. Using the NuMA-NG-P2A-Gal4 donor in combination with a UAS-H2B-Cherry reporter, we were able to document the dynamics of NuMA subcellular localization during mitosis (Figure 4D). In particular, in addition to the strong signal at spindle poles, we could observe the modest LGN-dependent cortical recruitment of NuMA during metaphase, and the LGN-independent increase in its cortical localization during anaphase and telophase that were previously described in vitro (Kotak et al., 2013). In our hands, this cortical localization of NuMA had so far been impossible to document in live conditions in the neuroepithelium with overexpression of fluorescently-tagged full-length NuMA (unpublished observations), illustrating the power of an endogenous tagging approach to document protein dynamic sub-cellular localization.

Finally, we illustrate the N-terminal tagging of Tubulin alpha1A1 (TUBA1A1) with a NeonGreen tag via the MMEJ approach. A short time-lapse sequence in a dividing neural progenitor shows that the endogenous tagging of alpha-tubulin offers a strong signal and an excellent signal/background ratio (Figure 4E).

Overall, these examples of endogenous protein tagging with fluorescent reporters illustrate that somatic knock-in is a powerful approach to document dynamic subcellular distribution of proteins of interest. With the MMEJ approach, which facilitates vector construction, it is also very rapid and simple to implement (see Discussion)

### Somatic knock-in for population-specific expression of reporters and lineage studies

In addition to protein tagging for live imaging, knock-ins allow to target specific cell populations based on the restricted expression of specific genes. Locus-specific expression of reporter genes has been used for decades in genetic models (eg. mouse, drosophila, zebrafish). In particular, binary systems using the Gal4/UAS transcriptional amplification system or the Cre/loxP indelible switch are systems of choice for the differential labelling and subsequent sorting of specific cell populations or for lineage tracing experiments (Buckingham and Meilhac, 2011). We evaluated the feasibility of similar experiments in the context of somatic knock-ins. We first targeted the Pax7 locus, which is only active in the dorsal half of the embryonic neural tube (Jostes et al., 1990). We generated a Pax7 HDR matrix introducing a pseudo-cleavage P2A sequence and the Gal4-VP16 transcription factor sequence at the C-terminus of Pax7. This matrix was coelectroporated with Cas9/gRNA vectors, together with a CAGGS-H2B-mRFP vector to report electroporation efficiency and an UAS-nls-EGFP vector to report the expression of Gal4. While the negative control Cas9/gRNA vector yielded virtually no GFP positive cells, Pax7 gRNAs led to a strong GFP fluorescence; on transverse sections, expression of GFP was restricted to the Pax7 domain in the dorsal neural tube, with virtually no GFP-positive cells in the ventral half, although the neural tube had been electroporated in the entire dorso-ventral axis, as revealed by the CAGGS-H2B-mRFP electroporation reporter. This indicates that knock-in of a Gal4-VP16 driver in target genes can be used to trigger strong and faithful cell-type specific expression from UAS responder elements. Similar results were obtained with a Pax7-P2A-Cre HDR donor vector, using a CAGGS-loxP-stop-loxP-mEGFP reporter instead of the UAS-nls-GFP in this case ((Morin et al., 2007); Figure 5).

**Figure 5.**
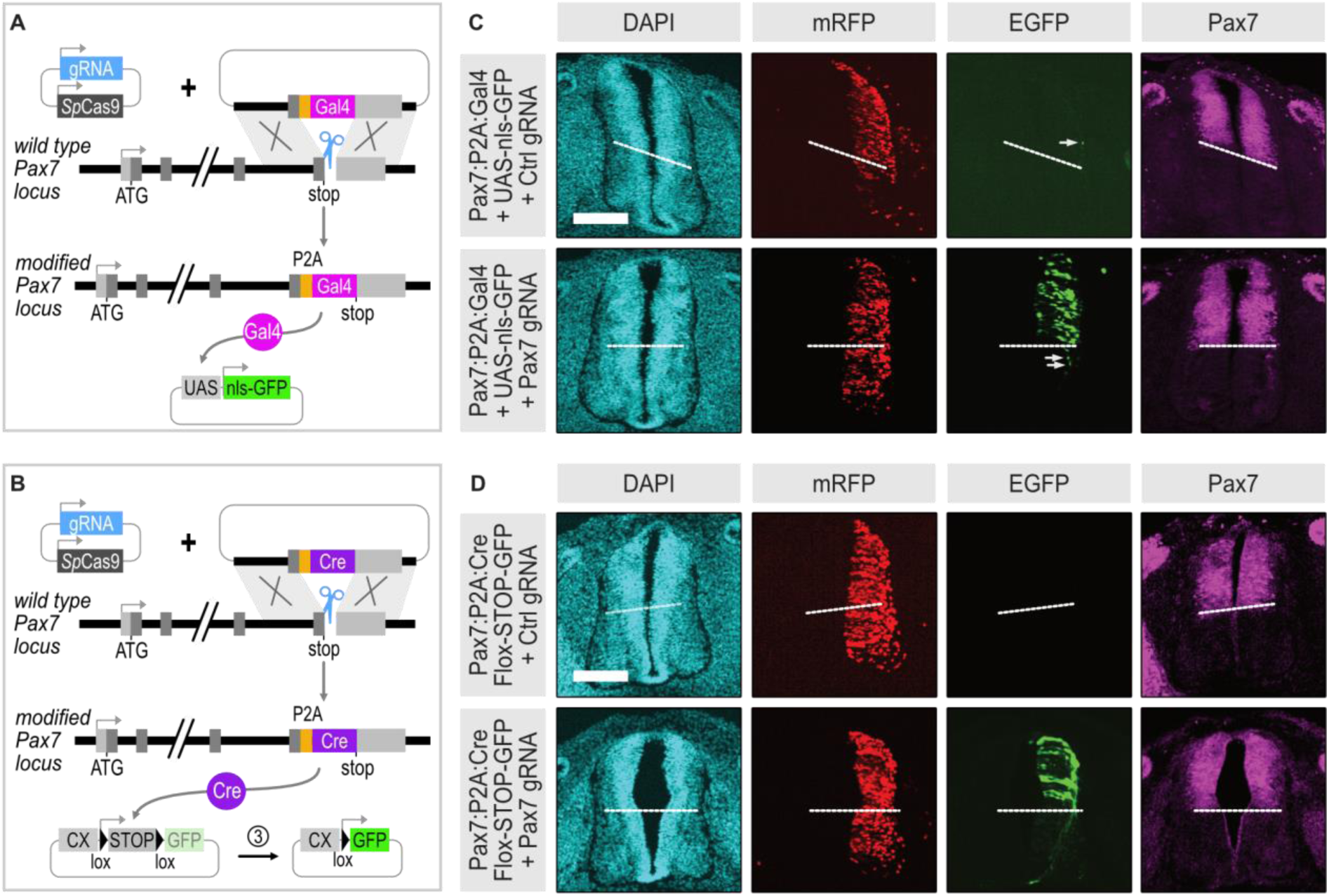
– Insertion of Gal4 and Cre recombinase for locus-specific binary activation of reporter genes. A-B. strategy for the insertion of the Gal4 transcriptional activator (A) or the Cre recombinase (B) at the Pax7 locus. The donor vector is designed to introduce the coding sequence of Gal4 or Cre as a fusion at the C-terminus. Introduction of a P2A pseudo-cleavage sequence between the last amino acid of the target gene and Gal4 or Cre ensures that the protein encoded at the target locus and Gal4 or Cre are present as separate proteins and perform their function independently. Gal4 or Cre are only expressed in the subpopulation of cell with a knock-in event in which the target gene is active, where they can drive expression from a responder plasmid. Gal4 directly activates transcription of a reporter via a UAS promoter (A). Cre excises a loxP flanked “stop” cassette to allow transcription of a GFP reporter driven by a ubiquitous CX promoter (B). C. Gal4 driven expression from a GFP responder plasmid upon insertion of Gal4 in the Pax7 locus. Embryos were electroporated at E2 with a mix of the Cas9 and gRNA expressing vector, a Gal4 donor vector, a CX-H2B-mRFP electroporation reporter, and a UAS-nls-GFP vector reporting the activity of the target locus in cells with successful knock-in events. Transverse cryosections were obtained at E4 at the thoracic level and immunostained with an anti-Pax7 antibody (magenta). Embryos with similar electroporation levels (as shown by the H2B-mRFP reporter, red) are displayed. Only rare GFP expressing cells are observed when a negative control gRNA is used (top row), while numerous cells are observed when the locus-specific gRNA is provided (bottom row), showing specificity, efficiency, and absence of background. Specificity is further demonstrated by the fact that GFP-positive cells are observed only in the Pax7-positive dorsal domain, revealed by the Pax7 antibody (magenta), with rare GFP positive cells outside of this domain (white arrows). While the Pax7 locus is switched off in differentiating dorsal neurons, the GFP reporter remains visible, due to the high stability of the GFP transcript and protein under Gal4 control. Scale bar, 100µm. D. Cre/loxP dependent expression of GFP upon insertion of the Cre recombinase in the Pax7 locus. Embryos were electroporated at E2 with a mix of the Cas9 and gRNA expressing vector, a Cre donor vector, a CX-H2B-mRFP electroporation reporter, and a CX-loxP-STOP-loxP-GFP vector reporting the activity of the target locus in cells with successful knock-in events. Transverse cryosections were obtained at E4 at the thoracic level and immunostained with an anti-Pax7 antibody (magenta). Scale bar, 100µm.

In the developing vertebrate spinal cord, neural progenitors initially amplify via symmetric (proliferative P-P) divisions, before switching progressively to neurogenic asymmetric (P-N) and symmetric terminal (N-N) divisions. The anti-proliferative transcription factor BTG2/Tis21 has been proposed to be absent from proliferative (P-P) progenitors, and specifically expressed in neurogenic (P-N and N-N) progenitors (Hämmerle et al., 2002; Iacopetti et al., 1999; Saade et al., 2013). While Tis21 expression dynamics is consistent with this model, appearing and increasing at the onset of neurogenic divisions, no lineage tracing experiments have been performed to confirm the identity of Tis21+ progenitors daughter cells with a cellular resolution in the spinal cord. We therefore sought to use the somatic KI approach to specifically label Tis21-positive progenitors and their progeny. We first constructed HDR donor vectors to target the insertion of P2A-Gal4 or P2A-Cre at the C-terminus of Tis21, and generated three gRNAs plasmids targeting Tis21. One of the gRNAs (gRNA2) yielded a very strong GFP signal when used in combination with the Tis21-P2A-Gal4 matrix and the UAS-nls-GFP reporter, whereas virtually no GFP-positive cells were observed when the control gRNA was used (Figure 6A-B). Similarly, we obtained a strong and specific signal when the Tis21-P2A-Cre donor and gRNA2 were used in combination with the CAGGS-loxP-polyA-loxP-EGFP reporter (Figure 6C-D).

**Figure 6.**
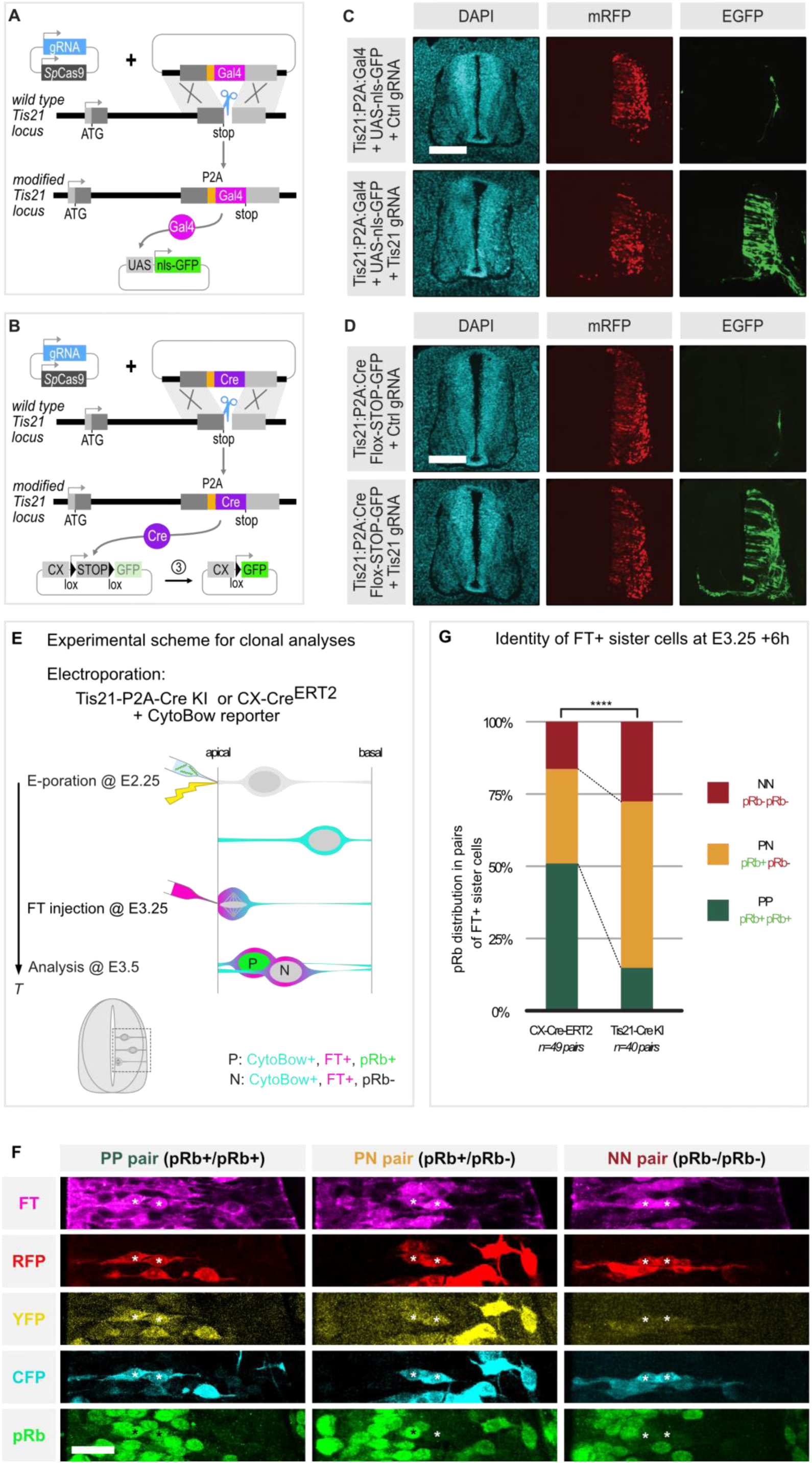
– Gene-specific lineage tracing via targeted insertion of a Cre recombinase. A-B. strategy for the insertion of the Gal4 transcriptional activator (A) or the Cre recombinase (B) at the C-terminus of Tis21. The donor vector is designed to introduce the coding sequence of Gal4 or Cre as a fusion at the C-terminus. Introduction of a P2A pseudo-cleavage sequence between the last amino acid of the target gene and Gal4 or Cre ensures that the protein encoded at the target locus and Gal4 or Cre are present as separate proteins and perform their function independently. Gal4 or Cre are only expressed in the subpopulation of cell with a knock-in event in which Tis21 is active, where they can drive expression from a responder plasmid. Gal4 directly activates transcription of a reporter via a UAS promoter. Cre excises a loxP flanked “stop” cassette to allow transcription of a GFP reporter driven by a ubiquitous CX promoter. C-D. Examples of Gal4 (C) or Cre (D) driven expression of GFP from the corresponding responder plasmids upon insertion of Gal4 or Cre in the Tis21 locus. Embryos were electroporated at E2 and transverse cryosections were obtained at E4 at the thoracic level. Only rare GFP expressing cells are observed when a negative control gRNA is used (top rows), while numerous cells are observed when the locus-specific gRNA is provided (bottom rows), showing specificity, efficiency, and absence of background. Scale bar, 100µm. E. Principle of the paired-cell clonal analysis. Embryos are co-electroporated at HH14 (yellow thunder) with either the Tis21-P2A-Cre donor and Tis21 Cas9/gRNA vector or with the Cx-Cre-ERT2 control plasmids, together with the CytoBow reporter. Embryos are injected with the FlashTag dye 24 hours after electroporation to label a synchronous cohort of mitotic progenitors, and collected six hours later. Anti-pRb and anti-GFP Immunofluorescence on thoracic vibratome sections determines the progenitor (pRb+) or prospective neuron (pRb-) status of FlashTag-positive electroporated sister cells, and allows to deduce the mode of division (P-P, P-N or N-N) of the mother cells. F. Representative two cell clone examples in transverse neural tube sections. From left to right panels: P-P, P-N and N-N pairs. Asterisks show pRb-positive progenitors (green) and pRb-negative future neurons in FlashTag-positive (magenta) pairs. The presence of an identical combination of CFP, YFP and RFP signals in pairs of FT-positive cells confirms their status of sister cells. Scale bar: 10µm G. Diagram indicating the percentage of P-P, P-N, and N-N clones in the Tis21-Cre and general progenitor populations at E3.5. P-P, P-N, and N-N stand for divisions producing two progenitors, one progenitor and one neuron, or two neurons, respectively. Chi-square test, ****p < 0.005.

In order to analyze the fate of Tis21-positive progenitors’ daughter cells, we introduced the Tis21-Cre matrix, Tis21-gRNA2 and Cas9 together with the MAGIC marker cytoplasmic reporter (Loulier et al., 2014). Upon Tis21 locus activation, cells with a KI event will express the Cre recombinase and recombine the MAGIC marker plasmids to activate a random combination of YFP, CFP and RFP reporters. The specific color combination activated in a progenitor will be inherited by its progeny, allowing for its clonal tracing based on color content. We then injected the far-red FlashTag dye in the neural tube 24 hours after electroporation (Figure 6E). FlashTag (FT) is a cell-permeant dye that is selectively incorporated by dividing cells in the neural tube during approximately half an hour after injection. It can therefore specifically label a cohort of cells undergoing mitosis at the time of injection (Baek et al., 2018; Telley et al., 2016). Embryos were harvested 6 hours after FT injection. To identify daughter cells fate, vibratome sections were immuno-stained with an anti-pRb antibody, which recognizes the hyper-phosphorylated form of the Rb protein. pRb is specifically detected in cycling progenitors after they pass the restriction point that occurs during G1, and until mitosis. Rb is not hyperphosphorylated in early G1, nor in cells that have exited the cell cycle. Hence, immunoreactivity for pRb becomes a discriminating factor for progenitors only after they have passed the restriction point. We have established that at E3.5, all cycling progenitors pass the restriction point within 6hrs after the previous mitosis (Mida et al, in preparation). Hence, 6 hours after FT injection, a FT+/pRb+ signal corresponds to a progenitor, while a FT+/pRb-signal corresponds to a post-mitotic neuron (Figure 6E). We analyzed pairs of FT-positive cells sharing a same MAGIC marker color combination (hence corresponding to pairs of sister cells born 6 hours before fixation from the mitosis of a Tis21-Cre positive progenitor) for Rb hyperphosphorylation. Pairs with two pRb+ cells result from a P-P (symmetric proliferative) division, while neurogenic divisions should contain only one (asymmetric, P-N) or zero (symmetric terminal, N-N) pRb+ cells (see characteristic examples in Figure 6F). As a control, we analyzed pairs of cells in the overall progenitor population at the same stage. Instead of the Tis21-Cre KI matrix and the Cas9-gRNA vector, we used a CAGGS-ERT2-Cre-ERT2 vector as an inducible source of Cre recombination (Matsuda and Cepko, 2007). 4OH-Tamoxifen was added to the embryos 18 hours after electroporation to trigger recombination of the MAGIC vectors in progenitors ahead of FT injection. FT was injected 24hae and embryos harvested 6hours after FT injection, as in the Tis21-Cre condition, corresponding to E3.5. At that stage, the control population contained roughly one half of proliferative P-P pairs (51 %, n= 25/49), and one half of neurogenic pairs consisting of 33% of asymmetric P-N pairs (n=16/49), and 16% of N-N pairs born from symmetric terminal divisions (n=8/49). In contrast the Tis21-Cre population yielded a majority (85%, n=34/40) of neurogenic divisions, consisting of 58% P-N divisions (n=23/40) and 27% N-N divisions (n=11/40). Symmetric proliferative P-P divisions were observed in 15% (n=6/40) of Tis21-Cre progenitors.

Overall, this analysis confirms a preferential expression of Tis21 in neurogenic progenitors, but also shows that Tis21 expression is not strictly restricted to this population (Figure 6G). More broadly, this set of experiments also demonstrates that the somatic knock-in of a Cre reporter can be used to selectively target and label a defined cellular population in order to characterize a specific parameter of this population (in the present case, the mode of division).

### Evaluation and suppression of background

When tagging proteins of interest with fluorescent reporters, we observed very little background signal (eg. signal in the presence of a control gRNA, or signal with aberrant subcellular localization in presence of a gene-specific gene). However, it has been suggested that the somatic insertion of reporters such as the Gal4 transcription factor or the Cre recombinase may lead to strong background signal due to a combination of low-level leaky episomal expression and high signal amplification inherent to these binary systems (Tsunekawa et al., 2016). In our initial tests described above, we were therefore pleased to find that the insertion of such cassettes at the Pax7 and Tis21 loci led to a specific reporter activation, with very few cells expressing the reporter when a control gRNA was used (Figures 5C and 5E and Figure 6B and 6D). However, in several other loci, we obtained a much stronger background. For example, Figure 7A illustrates that it was virtually impossible to distinguish between conditions using the control and specific gRNAs when we attempted to target the Gadd45g locus loci with the Cre recombinase. Similarly, very high background was observed when targeting the Notch1 locus with a Gal4 driver (not shown). The strong background signal in these cases presumably results from amplification of random, low-level leaky episomal expression of Gal4 and Cre driven by cryptic regulatory elements in the donor vector. These high background levels were observed with constructs using the large HDR architecture that we used when we started experimenting with somatic knock-ins. We reasoned that the MMEJ strategy, which relies on much shorter homology sequences, might be less susceptible to leakiness. Besides, the rapid linearization and the ensuing degradation of MMEJ vectors after electroporation might further suppress any residual leakiness. To test this hypothesis, we constructed a MMEJ donor vector that targets the Gadd45g with the Cre recombinase. Strikingly, this strategy efficiently and completely suppressed the background compared to the HDR version of the Gadd45g-P2A-Cre vector (Figure 7B). Similarly, we found that switching to a MMEJ architecture completely suppressed the low –but nonetheless detectable-background observed with the Pax7-Gal4 HDR construct presented above (Figure 7C-D).

**Figure 7.**
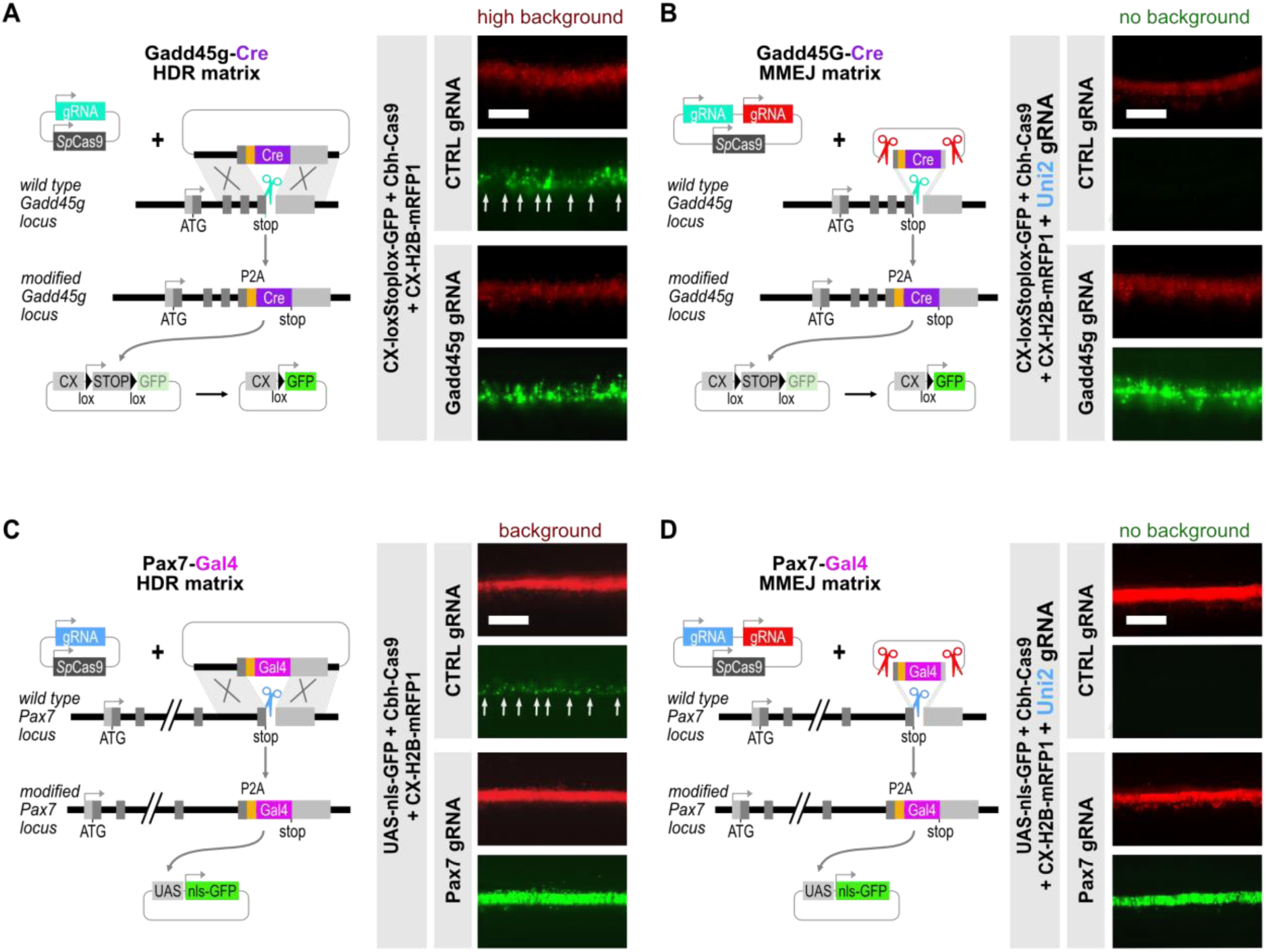
– MMEJ suppresses Gal4 and Cre background activity. A. The HDR approach using long arms of homology and circular vectors for the introduction of binary reporters may yield high background in some loci. Whole view of the electroporated neural tube 24 hours after electroporation of a mix of the Cas9/gRNA vector, a Gadd45g-P2A-Cre donor vector, a CX-H2B-mRFP electroporation reporter, and a Cx-lox-sto-lox-GFP vector reporting the activity of Gadd45g in cells with successful knock-in events. A strong background signal is observed in embryos having received the negative control gRNA (top panels), which are undistinguishable from embryos electroporated with the Gadd45g-specific gRNA. Scale bar: 300µm B. In contrast to the HDR strategy, the MMEJ approach targeting a Cre recombinase at the C-terminus of the Gadd45g locus yields no background in presence of the negative control gRNA (top panels), while the signal remains strong in embryos electroporated with the Gadd45g-specific gRNA. Scale bar: 300µm C-D: The HDR strategy targeting a Gal4 transgene to the Pax7 locus yields low but significant background in presence of the control gRNA (C, top panels); this background is entirely abolished when the MMEJ architecture is used to target the same locus (D, top panels), while the signal in presence of the Pax7 gRNA is similar between both strategies (bottom panels in C and D). Scale bars: 300µm

### Knock-in in other chick embryonic tissues

All the previous experiments were performed in the developing spinal neural tube, an embryonic structure which is easily accessible and ideal for in ovo electroporation. We also explored whether somatic knock-in can be achieved in other tissues that are more difficult to target. As a proof of principle, we used the highly expressed ACTB locus to ensure easy detection of knock-in events via direct fluorescence observation in the whole embryo. Figure 8A illustrates that knock-in of an ACTB-Cherry HDR matrix was achieved with high efficiency through ex ovo electroporation of unincubated embryos, as visualized 17 hours later in the epiblast. Figure 8B shows efficient knock-in of the ACTB-mNG HMEJ matrix in somites, 24 hours after electroporation at E2.5. In both cases, we did not observe any background in experiments using the negative control gRNA.

**Figure 8.**
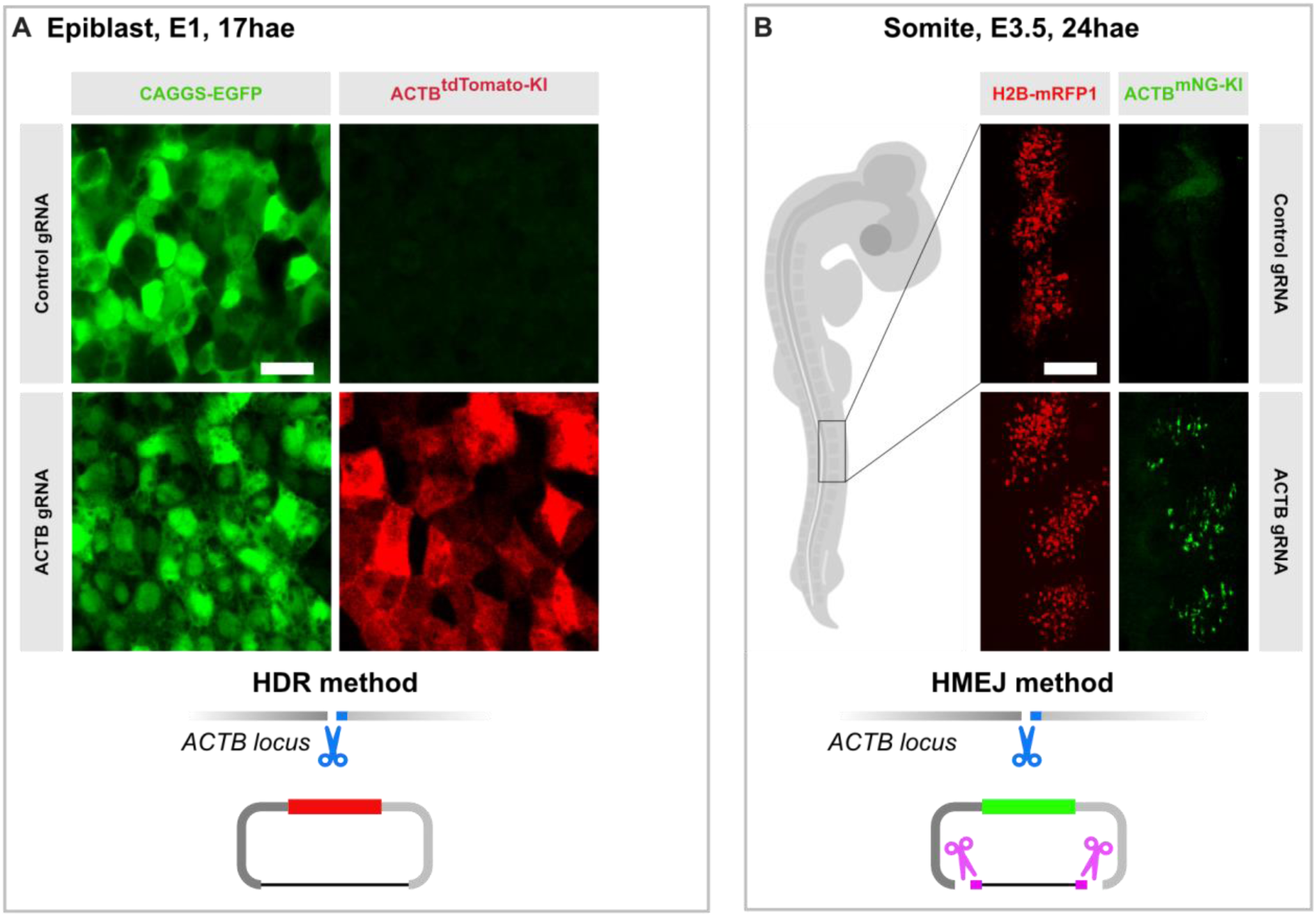
– somatic knock-in of fluorescent reporters in the chick epiblast and somites. A. A mix of ACTB single Cas9-gRNA vector, ACTB-Tomato HDR donor vector, and CX-H2B-GFP was electroporated ex-ovo in non-incubated embryos (HH stage 1), and electroporated embryos were cultured at 38°C for 17 hours before fixation and imaging. The images show a high magnification view of the epiblast that reveals a high frequency of ACTB-Tomato knock-in cells (red) in the electroporated population of cells (green) in presence of the ACTB gRNA, and absolutely no signal with the control gRNA. B. A mix of ACTB-uni2 dual Cas9-gRNA vector, ACTB-mNeonGreen HMEJ donor vector, and CX-H2B-mRFP was electroporated in the somatic compartment of E2.5 (HH stage 16) embryos, and the embryos were harvested one day later. The images show a high magnification view of 3 successive electroporated somites that reveals a high frequency of ACTB-mNG knock-in cells (green) in the electroporated population of cells (red) in an embryo electroporated with the ACTB-specific gRNA, and no knock-in events in the embryo electroporated with the control gRNA.

## Discussion

Our work compares several approaches for the implementation of somatic knock-ins in the developing avian embryo, primarily using the early spinal cord as a target tissue. We show that gene targeting with reporters is feasible, highly efficient, and easy to quickly implement via a strategy that requires minimal cloning steps. We also illustrate that the same strategy can be applied to other embryonic tissues.

### Implementation

Here, we used a plasmid-based approach, as opposed to the previously described method using a mix of ssDNA and protein/RNA approach used by Yamagata and Sanes in the embryonic chick retina (Yamagata et al., 2021). We found that the plasmid approach is reliable, easy to implement, and works better in our hands. In particular, the design and construction of MMEJ targeting vectors is very simple. The short homology means that no cloning or amplification of long genomic sequences is required. The entire arms of homology can be encoded in PCR oligonucleotides and constructs can be obtained in a single cloning step (supp figure 6). In our hands, the turnaround time from project design to in ovo validation can be less than two weeks, including the time required for oligonucleotides synthesis and delivery. One possible advantage of the riboprotein complex approach over plasmid-based methods is faster activity: in the former, all components are delivered “ready to use” in the cell upon electroporation, whereas in the latter Cas9 needs to be transcribed and translated, which imposes a delay of several hours after electroporation. However, we found that the plasmid-based approach resulted in knock-in events that were readily detected 18-24hae for all the loci that we have tested so far. We expect that this will be sufficient for most applications. Another possible caveat of the plasmid-based delivery is the longer persistence of Cas9 and gRNAs in cells, which may be a source of toxicity. In our hands, this has not been a problem: using Caspase 3 immunostainings on sections 24h after electroporation, we only observed few dying cells in the electroporated side of the neural tube. We did not quantify this effect, which appeared similar to what is routinely observed after electroporation of any plasmids in the neural tube.

### Efficiency

To evaluate knock-in efficiency, we used direct measurement in whole-mount embryos, and measured the ratio between cells with a knock-in event and cells expressing an electroporation reporter. We typically tested 2 or 3 gRNAs per locus, and obtained knock-in efficiency in the range of 10-40% with at least one of the tested guides for the 20 loci that we tested so far. We therefore anticipate that most genes can be efficiently tagged with this approach.

We note that the efficiency of knock-in appears to be higher in the neural tube than in the somite; in the spinal cord, it also appears to be highest when electroporation is performed at HH13-14, and to decrease in older embryos, although we did not attempt to quantify this in a systematic manner. This may be due in part to a lower electroporation efficiency at later stages or in the somite. Electroporation in the somite is notoriously less efficient than in the neural tube, and typically requires much higher DNA concentrations. In addition, it may depend on a differential regulation of DNA repair mechanisms between developmental stages and tissues. Strikingly, Keil and collaborators reported a decrease in the ratio of homologous to non-homologous repair pathways as neural progenitors switch from amplifying to neurogenic modes of division during mouse cortical development (Keil et al., 2020), which mirrors the observation that knock-in frequency in this tissue was much higher when in utero electroporation of knock-in components was performed at E12.5 than at E15.5 (Mikuni et al., 2016). Further work will be needed to optimize knock-in conditions in other developing tissues. In particular, a systematic comparison of HMEJ and MMEJ approaches may be useful, as the two approaches rely on different repair mechanisms for donor vector insertion. Several studies have reported that MMEJ knock-in efficiency can be improved in vitro by addition of exogenous components that favor the MMEJ mechanism over NHEJ, such as exonuclease 1 (Aida et al., 2016) or RB binding protein 8 (RBBP8-CtIP) (Nakade et al., 2018). We did not attempt to reproduce these results in ovo, as these approaches would require the addition of an additional vector in the electroporation mix, potentially mitigating the benefit by lowering the co-electroporation efficiency of all required components, in addition to potentially interfering genome wide with DNA repair mechanisms. However, a recent study showed that a fusion of the exonuclease1 to the C-terminus of Cas9 improves MMEJ targeting efficiency, reduces unwanted NHEJ at the targeted locus, and does not interfere detectably with DNA repair outside of the targeted locus (Shi et al., 2024). This approach might offer a way to further improve in ovo knock-in efficiency without increasing the number of components in the system.

### Targetability of loci

The knock-in insertion of fusion proteins at the N-or C-terminus of a protein of interest requires that gRNA target sequences are present in close proximity the start or stop codons. The ideal situation is that a gRNA target sequence overlaps the start or stop codons and is spread over the two arms of homology, as this ensures both that the cleavage will be close to the desired fusion site, and that the target sequence is not present in its entirety in one of the two arms of homology. In practice, this is not always possible. If the whole gRNA sequence is comprised in one arm of homology, a few bases should be switched, added or removed to “destroy” the recognition in the arm of homology and prevent targeting of the vector, and retargeting of the locus after recombination. For the MMEJ approach, in which short arms of homology are used, this means extending slightly the length of the arm. For N-terminal fusions, one should avoid gRNAs that are entirely comprised in the 5’ arm, as changing any bases upstream of the ATG might interfere with translation initiation. So far, including the examples presented in this study, we have targeted 20 different loci, either at the C-or the N-terminus (Supplementary Table 1A-B), and have found at least one gRNA yielding high knock-in efficiency for each of them.

### Fidelity and background

We found that there was very little background expression of reporters in conditions that use a control rather than a locus specific gRNA, even when using very sensitive amplification strategies based on the Gal4-UAS and Cre-loxP binary systems. In particular, the MMEJ strategy relying on short arms of homology and donor vector linearization had virtually zero background across 7 independent loci that we have tested so far (figure 5 and not shown). However, we cannot exclude that the approach induces off-target effects in the presence of locus-specific guides. Indeed, off-target activity of the Cas9/gRNA duplex could result in illegitimate insertions of the vector, leading to aberrant localization of the tag, or to deleterious effects for the cell. Should they occur, both types of effects would in principle be guide specific, and can therefore be identified and circumvented by testing at least two different gRNAs per target gene. We anticipate that the risk of illegitimate off-target insertions leading to detectable reporter signal will generally be low. However, off-target activities may also alter cell function and developmental events, and therefore indirectly affect reporter expression or localization resulting from a legitimate integration event. Overall, selecting gRNAs with a low off-target prediction score, using dedicated algorithms such as the one proposed by the CRISPOR platform that we used for our gRNA selection, and testing at least two independent gRNAs is recommended, especially in experiments that would combine a knock-in strategy with orthogonal functional approaches (eg. gain and loss of function) to monitor and compare a protein’s subcellular distribution between different experimental conditions.

### Knock-ins for the documentation of subcellular localization and dynamics

In contrast to fluorescent reporter insertions in the ACTB and TUBA1A1 loci, for which the fluorescent signal was visible by direct inspection of embryos directly in the egg with a fluorescence stereomicroscope 18 to24 hours post electroporation, this was not the case for any of the other tagged proteins described in this study. This is consistent with our transcriptomic data showing that these genes are expressed on average at 10-50x lower levels compared to ACTB in the neural tube. All these fusion proteins, as well as several others not described here, were nonetheless detected in live and fixed confocal microscopy at high resolution. Overall, endogenous tagging appears as a superior strategy to monitor subcellular dynamics, as it suppresses overexpression and fixation artefacts. To overcome low expression levels, the use of very bright fluorescent reporters such as NeonGreen is highly recommended, especially for live imaging. In addition, tandem repetitions of the reporter can be used to increase signal intensity. We have not attempted to systematically quantify whether increasing the size of the inserted sequence reduces the frequency of knock-in events, but we routinely use tandem copies of NeonGreen, or NeonGreen-P2A-Gal4 cassettes, and have not observed that this strongly affected the efficiency of the approach. Indeed, we have successfully inserted long fragments of up to 4.5kb (not shown).

### HMEJ versus MMEJ?

Although HMEJ, using the linearization of a vector with long arms of homology, resulted in higher efficiency than MMEJ at the ACTB locus, and might be a good choice for dual gene tagging strategy, our favored approach is currently the MMEJ, thanks to the easy implementation of the cloning strategy. MMEJ is also probably a better approach for N-terminal fusions, as the short arms of homology avoid the introduction of regulatory sequences in the left arm of the donor vector. We would therefore recommend starting with a MMEJ strategy, which should be sufficient for most applications, and possibly switching to a HMEJ approach for specific applications where the highest knock-in frequency is necessary, such as multiplex gene targeting. As mentioned above, MMEJ efficiency might be further improved by implementation of the Cas9-Exonuclease1 strategy (Shi et al., 2024).

### Future applications in lineage and functional studies

We demonstrate here that the somatic knock-in approach offers an efficient and specific way to target and label specific cell populations with Gal4 and Cre drivers. This opens the way to refined lineage studies and the measurement of parameters in specific cell subpopulations. We have shown here that inserting a Cre or Gal4 recombinase at the Tis21 locus allows to access and label a population of cells enriched in neurogenic progenitors, and particularly in asymmetrically dividing progenitors. We illustrate this application to characterize new parameters of asymmetric cell division in a separate study (Bunel et al, in preparation). In addition to reporters or Gal4 or Cre drivers, somatic knock-ins can be used to directly place the coding sequence of a gene of interest under the control of the regulatory sequences of another locus. This allows to achieve an exogenous expression that is reproducibly controlled in time, space, and level, and opens the way to refined functional studies that avoid the caveats of massive and ubiquitous overexpression. We also illustrate this application in a parallel study, where we used specific insertion of the cell cycle regulator CDKN1c/p57kip2 in the Pax7 locus to achieve a low-level expression restricted to dorsal progenitors (Mida et al, in preparation). Considering the ease of deployment of such strategies, which do not necessitate the establishment of transgenic lines and complex crossing schemes, we anticipate many more applications of this strategy, such as the targeted expression of modifying enzymes for the cell-type specific exploration of chromatin conformation or transcription factor binding sites (Van Den Ameele et al., 2019).

## Conclusion

With a turnaround time of less than two weeks from the design of a knock-in strategy to its in ovo validation by electroporation in the chick embryo, the MMEJ approach for somatic knock-ins is arguably the fastest route to document the expression pattern and the subcellular localization and dynamics of any protein of interest in the avian embryo. It will also be a powerful tool for the development of lineage and functional studies based on conditional systems. Our study has mostly focused on the spinal cord neural tube, which is one of the most easily accessible tissue in the embryo. However, we show that the tools presented here are directly transferrable to other embryonic tissues in the chick embryo. The same designs should also work in other avian and in mammalian species. Somatic CRISPR/Cas9 based knock-ins represent a simple and powerful approach for cell and developmental in ovo studies in birds, that brings the power of modern genetic approaches to a historical developmental biology model while circumventing the long and cumbersome process of establishment of stable transgenic lines.

## Materials and Methods

### Plasmids

#### gRNA and Cas9 expression vectors

We used the X330 vector (http://www.addgene.org/42230) described in (Cong et al., 2013) for the simultaneous expression of the Cas9 nuclease and gRNA. Each gRNA vector was constructed by cloning a duplex of oligonucleotides in BpiI-digested X330. For gRNAs starting with a G, oligonucleotides were chosen as follows: 5’-CACC[20bpsensegRNA]-3’ and 5’-AAAC[20bpantisensegRNA]-3’; for gRNAs starting with A, C, or T, a G must be added upstream of the gRNA sequence to initiate translation from the U6 RNA polymerase. Oligonucleotides were therefore designed as follows: 5’-CACCG[20bpsensegRNA]-3’ and 5’-AAAC[20bpantisensegRNA]C-3’.

For double gRNA constructs, the uni2 gRNA target sequence (GGGAGGCGTTCGGGCCACAG; (Welker et al., 2021) was first introduced into the X330 vector to generate the uni2 gRNA plasmid (X-1242); the “empty” U6-gRNA cassette was then PCR amplified from X330 with U6-fw 5’-*Aattctgcagacaaatggct*GAGGGCCTATTTCCCATGAT-3’ and gRNACass-rev 5’-Cgtaagttatgtaacgggtacctcta-3’ and introduced into XbaI-digested X-1242 by Gibson assembly cloning to generate the dual gRNA cassette dgRNA-empty-uni2 vector (X-1251). Locus-specific gRNAs were then introduced in the empty gRNA cassette in BpiI-digested X-1251 as described above.

The targeted genomic sequences of genes tested in this study and the associated gRNAs are available in Supplementary Table 1A.

#### Donor vectors

Knock-in matrices were generated via Gibson assembly or conventional restriction digestion and ligation cloning. For HDR vectors, the long arms of homology were PCR amplified from chick genomic DNA. Several GC rich fragments could not be amplified by PCR, and were therefore synthetized (Eurofins). All other elements (fluorescent reporters, Gal4-VP16, Cre recombinase, short arms of homology) were PCR amplified from plasmid DNA.

MMEJ vectors are based on a 1.8kb minimal backbone, consisting of an AmpR antibiotic resistance gene, and an ori2 origin of replication, derived from the pUC19 backbone. They are routinely constructed in a single cloning step via Gibson cloning of two PCR fragments (supplementary figure 5). In short, one PCR fragment (the “backbone fragment”) spans the vector backbone (AmpR and ori2) flanked by Uni2 gRNA target sites and part of the arms of homology; the other PCR fragment (the “insert fragment”) contains the desired reporter, flanked by a complementary part of the arms of homology.

This double PCR method allows to keep oligonucleotide length below 50 bases, and to use the minimal synthesis scale (0.01µmol) from our usual manufacturer (Eurofins genomics).

The insert fragment is amplified with oligos 1 and 2 designed as follows:

1: 5’-[<u>sense_left_arm</u>-reporter-Nter]-3’ in forward orientation

2: 5’-[<u>antisense_right_arm</u>-reporter-Cter]-3’ in reverse orientation

The backbone fragment can be amplified with oligos 3 and 4 from any existing linearisable MMEJ or HMEJ vector in which AmpR and ori2 are already flanked by Uni2 target sites, using oligos recognizing Uni2 flanked on their 5’ end with the desired portion of the arms of homology (oligos 3 and 4 in figure). As Uni2 sites are immediately flanked by XbaI (TCTAGA) and EcoRI (GAATTC) sites, 2 specific bases (GA or TC) are added downstream of the uni2 sequence on the 3’ end of oligos 3 and 4 to provide strand specificity and prevent them from annealing on the same uni2 sequence.

Oligos 3 and 4 are designed as follows:

3: 5’-[antisense_left_arm]-CCKCTGTGGCCCGAACGCCTCCCtc-3’

4: 5’-[sense_right_arm]-CCKCTGTGGCCCGAACGCCTCCCga-3’

CCKCTGTGGCCCGAACGCCTCCC corresponds to the uni2 target site (GGGAGGCGTTCGGGCCACAGMGG) in reverse orientation where MGG corresponds to the PAM (either AGG or CGG, which we have both tested in MMEJ vectors).

The 5’ tails of oligos 1, 2, 3 and 4 are designed to be reverse complementary (1 with 3, 2 with 4) over a minimum of 15 bases to generate a 35bp arm of homology on both sides of the reporter via Gibson recombination.

This double PCR method allows to keep oligonucleotide length below 50 bases, allowing rapid production at the minimal synthesis scale (0.01µmol) from our usual manufacturer (Eurofins genomics).

In routine, we prefer to amplify the backbone fragment from our existing linearisable vector X-1308 (a 4.6kb long HMEJ vector targeting Cherry to the ACTB locus with long arms of homology) rather than from shorter MMEJ vectors (typically 2.5kb long). We find that this limits the risk of contamination with the supercoiled matrix during gel purification of the 1.87bp PCR fragment, and removes the need from a DpnI digestion step.

For the identification of positive clones, we use the following oligonucleotides (5 and 6 on figure): Amp-rev (ATAGGGGTTCCGCGCACA) and Ori-fw (Acgcggcctttttacggttc) and perform PCR directly on bacterial colonies. 5µl of PCR reaction is loaded on gel to check the presence of a band of the desired size. For positive clones, the remaining PCR reaction is purified on columns and sent for sequencing with Amp-Rev and ori-fw oligos. In principle, for reporters shorter than 800bp (eg most fluorescent proteins or Gal4), sequencing from 1 end should be sufficient to obtain a complete sequence of the whole uni2-left arm-reporter-right arm-uni2 cassette. However, we have noticed that the uni2 sequence is subject to frequent sequencing errors when read at the 3’ end of a sequencing reaction, and therefore recommend systematic sequencing from both ends. We do not sequence the PCR amplified backbone, as the presence of a colony implies that it has functional Amp resistance and origin of replication.

### Linear donor DNA

For experiments using the ssDNA ACTB-GFP11 donor oligonucleotide (Supp Figure 1), a ssODN was purchased from IDT, resuspended in IDT’s Nuclease Free Duplex Buffer (30 mM HEPES, pH 7.5; 100 mM potassium acetate) at a concentration of 80µM, and added to the electroporation mix at a final concentration of 16µM. The sequence of the ACTB-GFP11 ssODN is as follows: 5’-CCAGCAGATGTGGATCAGCAAGCAGGAGTACGATGAATCCGGACCCTCCATTGTCCACCGCAAATGCTTCGGTGGCGGCcgtgaccacatggtccttcatgagtatgtaaatgctgctgggattacaTAAGACTGTTACCAACACCCACACCCCTGTGATGAAACAAAACCCATAAATGCGCATAAAACAAGACGAGATT-3’.

For experiments using double stranded ACTB-NeonGreen PCR donor sequences, 5’-biotinylated oligonucleotides (Biot-ACTB-fw: 5’-AATCCGGACCCTCCATTGTCC-3’ and Biot-ACTB-rev 5’-CATCACAGGGGTGTGGGTGTT-3’) designed to amplify a 800bp fragment corresponding to the NeonGreen sequence flanked by 35bp long arms of homology from the ACTB-NG HDR vector (X1027) were purchased from Eurofins. After amplification with the DreamTaq PCR mix (ThermoFischer), the PCR product was purified on columns (Promega Wizard purification kit) and added to the electroporation mix at a concentration of 0.133µg/µl.

### Fertilized eggs

JA57 chicken fertilized eggs were provided by EARL Morizeau (8 rue du Moulin, 28190 Dangers, France) and incubated at 38°C in a Sanyo MIR-253 incubator. Embryos used in this study were between E2 (HH14) and E3 (HH14+24 h). The sex of the embryos was not determined. Under current European Union regulations, experiments on avian embryos between 2 and 4 days in ovo are not subject to restrictions.

### In ovo and ex ovo electroporation

Electroporations in the neural tube were performed at HH13-14 by applying 5 pulses of 25V for 50ms, with 100ms in between pulses. Electroporations were performed using a square wave electroporator (NepaGene CUY21SC Square Wave Electroporator, or BTX ECM-830 Electro Square Porator, or Ovodyne Intracell TSS20) and a pair of 5 mm Gold plated electrodes (BTX Genetrode model 512) separated by a 4 mm interval.

In most cases, and unless specified otherwise in figure legends, donor vectors and Cas9/gRNA vectors were used at a concentration of 0.8µg/µl each. Electroporation reporters and UAS reporters were used at concentrations ranging from 0.1 to 0.4µg/µl.

Ribonucleoprotein complexes were assembled as follows: 20 nt control and ACTB#3 target sequences were used to design 36 base long single stranded crRNA sequences. crRNA oligonucleotides were purchased from Integrated DNA Technologies (IDT) and resuspended in IDT’s Nuclease free duplex buffer at a concentration of 100µM. crRNA were mixed at equimolar concentration with a tracrRNA (purchased from IDT: ALT-R and resuspended in the same buffer) to obtain a 50µM trRNA mix in 10µl aliquots, and annealed by heating 5 min to 95C and cooling down to RT. Purified Cas9 protein (30µM in 10mM HEPES, 150mM KCl, a gift from A. De Cian and J.P. Concordet, was mixed 1:1 (vol:vol) with the trRNA duplex and incubated for 20 min at 37C to promote complex formation. 1µl of this mix was then complemented with Fast Green and a DNA reporter plasmid (pCX-H2B-mRFP) and the donor fragment consisting of either a single stranded DNA oligonucleotide or a double stranded PCR fragment, in a total volume of 5µl. Final concentrations in the mix are Cas9 protein 3µM, trRNA duplex 5µM, reporter plasmid 0.1µg/µl, and ssODN or PCR fragments at the concentration indicated above.

In ovo electroporation in the somite was performed essentially as described in Scaal et al (Scaal et al., 2004). Donor and Cas9/gRNA vectors were used at final concentrations of 2µg/µl each, and the CX-H2B-mRFP reporter was used at a concentration of 0.4µg/µl.

Ex ovo electroporation of unincubated chicken embryos: stage 1 embryos were collected on a filter paper ring and electroporated in a custom-made electroporation chamber using the Electroporator NEPA21 type II (NepaGene), with two 5ms poring pulses of 15V, 50ms delay, and three 50ms transfer pulses of 10V, 500ms delay. Plasmids were electroporated at a concentration of 1µg/µl. Embryos were then cultured ex ovo for 17h on a semisolid nutritive medium containing a mix of albumen (50%), agarose (0,2%), glucose and NaCl in a humidified chamber. Electroporated embryos were then fixed in 4% formaldehyde and imaged using a Zeiss LSM 900 confocal microscope.

### Visual evaluation of knock-in efficiency

For all our constructs tested in the neural tube, we performed a visual evaluation of knock-in efficiency using a Leica MF205 fluorescence dissection binocular. Images of electroporated neural tubes were acquired using a monochrome camera (PointGrey BFLY-U3-23S6M-C), and micromanager software. For consistency, we used the same 80x magnification factor on the dissection scope and a 2x ROI in micromanager for all images presented in this study. Images were taken either directly in the live embryo in the egg, or after dissection, removal of extraembryonic membrane and “pinning” of the embryo face down with 0.2mm iron minutiae on a India-ink tinted sylgard dish. In these cases, images were acquired either directly from unfixed embryos in PBS or after 1-hour fixation in 4% Formaldehyde in PBS and 3 PBS washes.

### Imaging

Confocal Images were acquired on an inverted microscope (Nikon Ti Eclipse) equipped with a spinning disk confocal head (Yokogawa CSUW1) with Borealis system (Andor) and a sCMOS Camera (Orca Flash4LT, Hamamatsu) and the following Nikon objectives: 10x (CFI Plan APO λ, NA 0.45), 20x (CFI Plan APO λ, NA 0.75), 40x (CFI Plan APO λ, NA 0.95) and 100x (CFI APO VC, NA 1.4, oil immersion). The setup is driven by MicroManager software (Edelstein et al., 2010) and equipped with a heating enclosure (DigitalPixel, UK) for live imaging.

### Quantification of Insertions with CellPose2

To quantify the number of single and double knock-in insertions, we acquired en-face images from flat mounted neural tubes from fixed embryos. We acquired 1µm spaced z stacks (3 to 7 µm full depth) at the level of the apical surface with a 100x objective. We used 4 to 6 embryos per condition, and for each embryo, we acquired a minimum of 4 apical fields of at least 65µmx65µm (this corresponds to the standard ROI crop value on our setup). For each field, we generated a z-projection of 3 z-levels centered on the maximal ZO1 signal intensity, and saved an individual tiff file for each channel. We loaded the ZO1 image in Cellpose2 (Pachitariu and Stringer, 2022), used the automated cell diameter calibration, and directly used the preloaded Cyto2 model to count the number of cells (attempts to train a custom model with our iRFP-ZO1 or ACTB-knock-in images did not improve the quality of segmentation). For each image, we corrected the output by counting the few ZO1-positive cells (generally less than 3% of the total count) that had not been detected by cellpose2 and adding then to the total count to obtain the number of electroporated cells in the field. We then processed the corresponding ACTB-NG and ACTB-Cherry images (either NeonGreen for single knock-in, or both NeonGreen and Cherry for double knock-in) in the same way to calculate the number of knock-in events in the field. knock-in efficiency is calculated as follows:

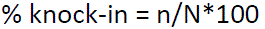

Where n is the number of cells with a knock-in event and N is the number of electroporated cells (ZO1-positive) counted in cellpose2 in the same field.

For double color knock-in experiments,

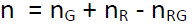

where n_G_ and n_R_ are respectively the numbers of knock-in cells counted in the green and red channels in cellpose2, and n_RG_ is the number of double-positive (red and green) cells counted by eye in the original multicolor image opened in Fiji.

For double Mib1-NG and ACTB-Cherry experiments, the number of iRFP-ZO1 electroporated cells and ACTB-Cherry knock-in cells in each image were counted in Cellpose2 as described above. Cellpose2 could not segment efficiently the Mib1-GFP signal, so the number of Mib1-GFP positive and of double Mib1-GFP and ACTB-Cherry positive cells were counted manually counted in Fiji.

### En-face culture for live imaging of the neuroepithelium

En-face culture of the embryonic neuroepithelium was performed at E3 (24 h after electroporation). After extraction from the egg and removal of extraembryonic membranes in PBS, embryos were transferred to 37 °C F12 medium and pinned down with dissection needles at the level of the hindbrain in a 60 mm diameter Sylgard dissection dish. A dissection needle was used to slit the roof plate and separate the neural tube from the somites from hindbrain to caudal end on both sides of the embryo. The neural tube and notochord were then transferred in a ∼50µl drop of F12 medium to a glass-bottom culture dish (MatTek, P35G-0-14-C) and medium was replaced with 1ml of 1% low melting point agarose/F12 medium stabilized at 38°C. Excess medium was quickly removed so that the neural tube would flatten with its apical surface facing the bottom of the dish, in an inverted open book conformation. After 30s of polymerization on ice, an extra layer of agarose medium (200 µl) was carefully added to embed the whole tissue and left to harden for 30 additional seconds. 2-3 mL of 38°C culture medium was added (F12/1mM Sodium pyruvate) and the culture dish was transferred to the 38°C chamber of the spinning disk confocal microscope.

## Authors contribution

Conceptualization: X.M.; Formal analysis: X.M., E.F.; Funding acquisition: X.M., M.M.; Investigation: A.P.-V., B.M., R.G., O.A.-P., B.D., E.F., S.T., J.G., M.M., X.M.; Methodology: X.M.; Project administration: X.M.; Supervision: X.M.; Validation: X.M.; Visualization: X.M.; Writing – original draft: X.M.

## Acknowledgements

We thank A. De Cian and J.P. Concordet for the generous sharing of purified Cas9 protein. We acknowledge the generous help from the labex MemoLife which allowed the initiation of this project through a collaborative grant to the Morin and Manceau labs. Work in the Morin lab was additionally supported by grants from the Fondation pour la Recherche Medicale (FRM EQU202003010547) and the Agence Nationale pour la Recherche (SYMASYM ANR-18-CE16-0021-01). B. Mida was supported by doctoral grants from the French Ministry of Higher Education and Research (MESR) and the Labex MEMO LIFE. This work has received support under the program « Investissements d’Avenir » launched by the French Government and implemented by the ANR, with the reference ANR-10-LABX-54 MEMO LIFE. The authors declare no competing financial interests.

## Figures and legends

**Supplementary Figure 1.**
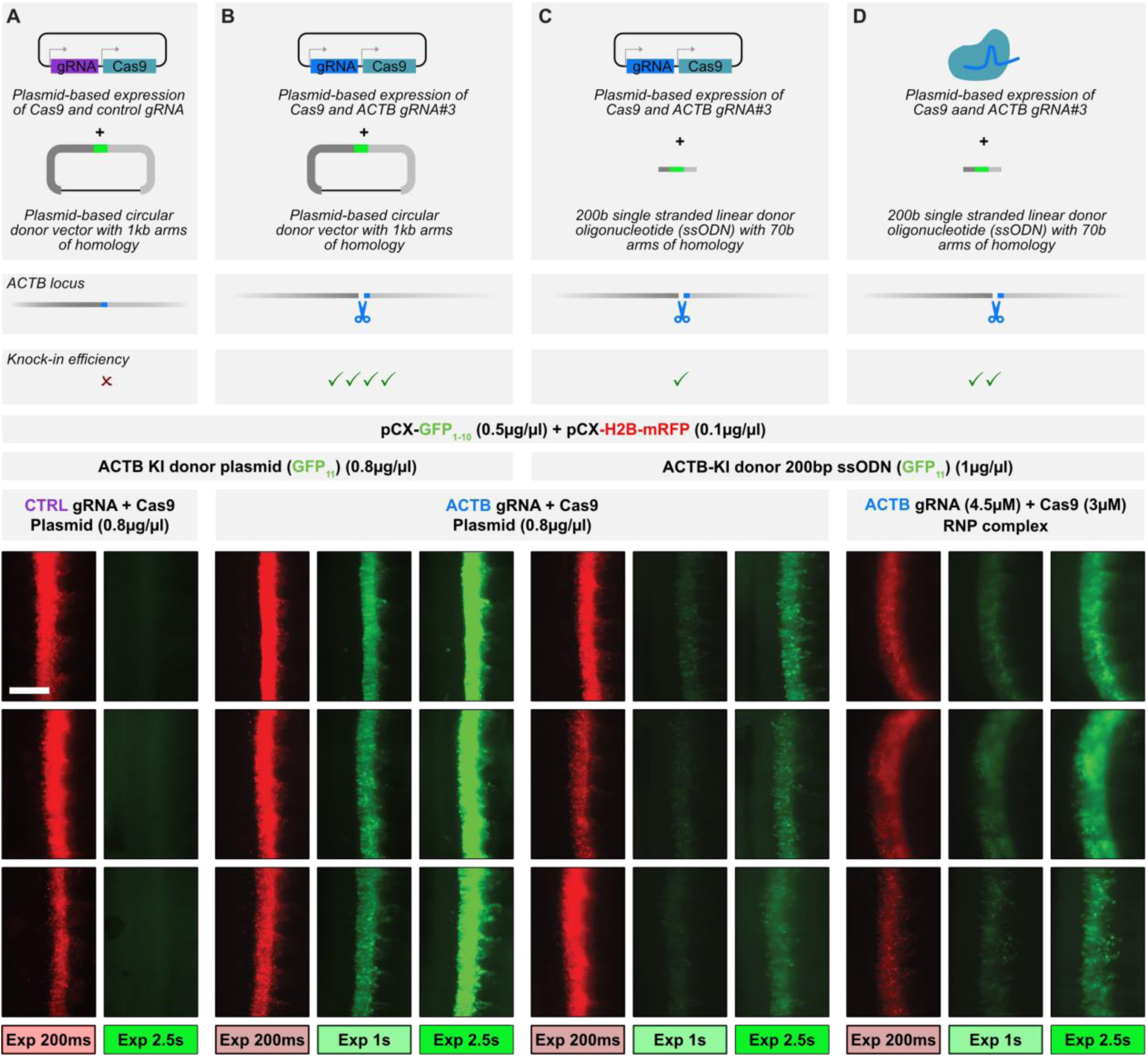
– Comparison of strategies based on plasmid-encoded or in vitro preassembled ribonucleoprotein Cas9/gRNA complexes for the insertion of short tags. Schemes in the top row of panels A to D represent the targeting strategy used in the different conditions. Scale bar, 300µm. A-B. A homologous recombination donor vector targeting a split GFP_11_ tag at the C-terminus of the ACTB locus was electroporated in the neural tube of stage 14 embryos with a Cas9-gRNA vector expressing the Cas9 protein and either a negative control gRNA (A) or gRNA#3 targeting the C-terminal region of ACTB (B). An electroporation reporter (CX-H2B-mRFP) and a vector expressing the complementary moiety of split GFP (CX-GFP_1-10_) were coelectroporated. Embryos were photographed directly in the egg 24 hours after electroporation. 3 representative embryos with similar electroporation level (red signal) are displayed for each condition. As expected, no KI events are observed in the negative control, even with long (2.5s) exposure, whereas a strong signal is observed with the ACTB gRNA. C. In condition C, the donor vector was replaced with a 200 bases long single stranded oligonucleotide in which the GFP_11_ tag sequence is flanked by ∼70 bases long sequences of homology to the ACTB locus. In these conditions, a modest GFP signal is observed, the intensity is much weaker than in condition B. D. In condition D, the donor vector was replaced with a 200 bases long single stranded oligonucleotide in which the GFP_11_ tag sequence is flanked by ∼70 bases long sequences of homology to the ACTB locus, and the Cas9 and gRNA were introduced as an in vitro preassembled ribonucleoprotein complex. In these conditions, the GFP signal is stronger than in condition C, but the intensity remains weaker than in condition B. This is probably due to low electroporation efficiency of the solution containing a mix of DNA and ribonucleoprotein complex, rather than low knock-in efficiency, as the electroporation reporter signal (red) is also consistently weaker in condition D compared to conditions A to C, in which the electroporated solutions only contain DNA.

**Supplementary Figure 2.**
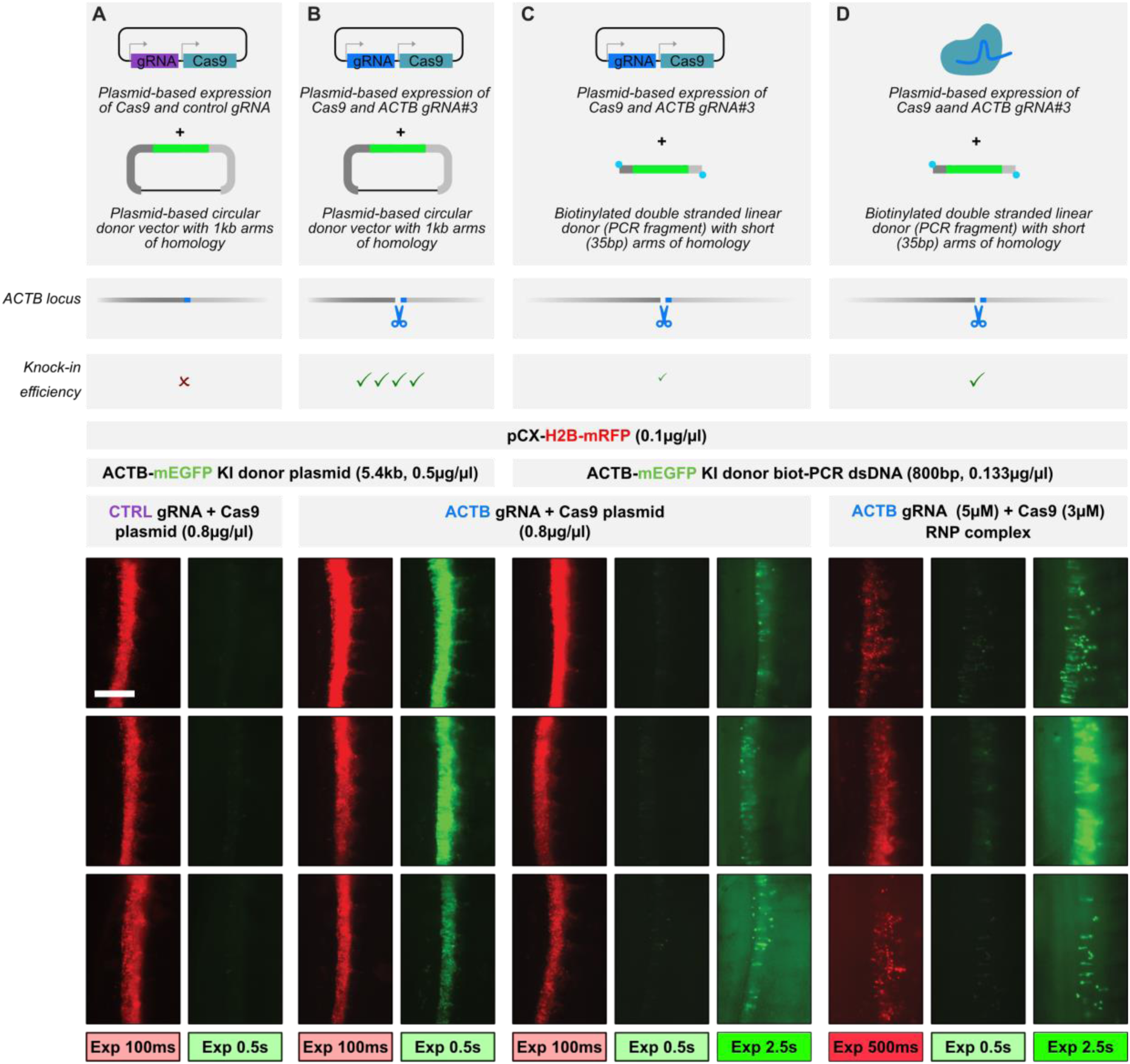
– Comparison of strategies based on plasmid-encoded or in vitro preassembled ribonucleoprotein Cas9/gRNA complexes for the insertion of long tags. Schemes in the top row of panels A to D represent the targeting strategy used in the different conditions. Scale bar, 300µm. A-B. A homologous recombination donor vector targeting a EGFP at the C-terminus of the ACTB locus was electroporated in the neural tube of stage 14 embryos with a Cas9-gRNA vector expressing the Cas9 protein and either a negative control gRNA (A) or gRNA#3 targeting the C-terminal region of ACTB (B). An electroporation reporter (CX-H2B-mRFP) was coelectroporated. Embryos were photographed directly in the egg 24 hours after electroporation. 3 representative embryos with similar electroporation level (red signal) are displayed for each condition. As expected, no KI events are observed in the negative control, whereas a strong signal is observed with the ACTB gRNA. C. In condition C, the donor vector was replaced with a PCR-generated single stranded linear fragment in which the EGFP sequence is flanked by ∼35 bases long sequences of homology to the ACTB locus. The PCR fragment was generated with 5’-biotinylated primers. In these conditions, a very modest GFP signal is observed, and is only detected when using very long exposure. D. In condition D, the donor vector was replaced with a biotinylated PCR-generated single stranded linear fragment in which the EGFP sequence is flanked by ∼35 bases long sequences of homology to the ACTB locus, and the Cas9 and gRNA were introduced as an in vitro preassembled ribonucleoprotein complex. In these conditions, very few KI events are detected and the GFP signal is extremely weak and barely detectable with short exposure time. We note that the electroporation reporter signal (red) is consistently weaker in condition D compared to conditions A to C, (the images were acquired with 500ms exposure in the red channel for this condition, compared to 100ms for conditions A-C), probably reflecting low electroporation efficiency of the solution containing a mix of DNA and ribonucleoprotein complex compared to electroporated solutions in conditions A-C which only contain DNA.

**Supplementary Figure 3.**
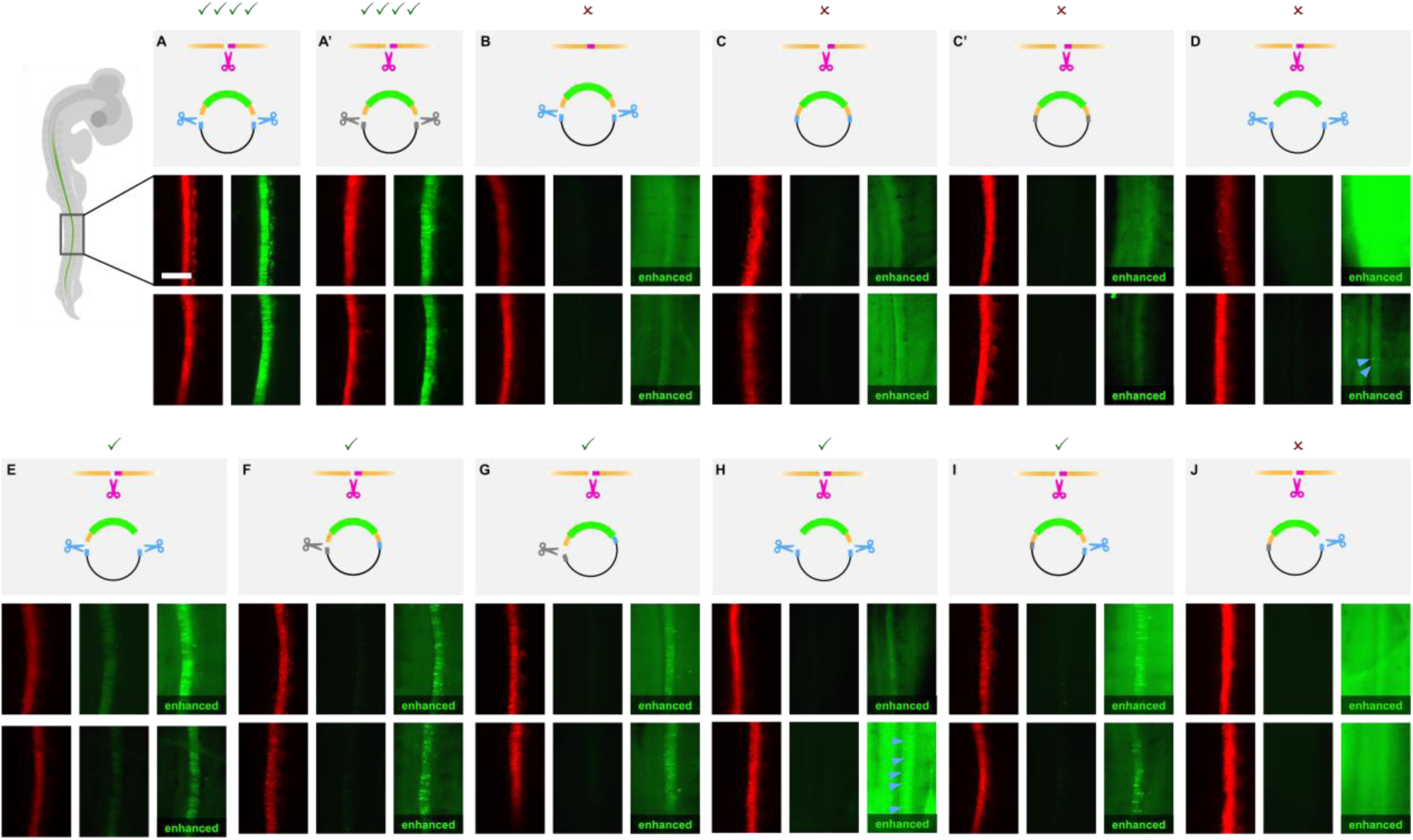
– MMEJ requires two donor arms each flanked by a linearization site for efficient knock-in production. We evaluated the requirement for arms of homology and linearization for efficient knock-in in the MMEJ strategy. We generated a series of mNeonGreen donor vectors targeting the ACTB with different combinations of presence or absence of the left and right arms of homology and of presence or absence of the left and right linearization sites. Panels A-J each show a different condition. In each panel, the characteristics of the vector are depicted schematically in the top row, and two representative examples with similar electroporation quality (as judged from the intensity of the signal from a CX-H2B-mRFP electroporation reporter in the red channel) are shown for each condition. In all conditions, except condition B, a gRNA targeting the ACTB locus was used (pink scissors in top schematic representations). Green color in the vector represents the mNeonGreen sequence, orange flanking regions correspond to left and right arms of homology, and blue lines indicate the presence of the uni2 linearization sites. In addition to uni2, we used an alternative linearization site (gray lines in vector schemes) which is recognized by another gRNA (uni1 gRNA). The presence of blue or gray scissors next to the donor vector scheme indicates that a Cas9/gRNA vector expressing the corresponding gRNA (uni2 or uni1 gRNA) was included in the electroporation mix of the corresponding condition. Only conditions A and A’, in which the donor vector contains a left and a right arm of homology and two linearization sequences, yield strong NeonGreen signal that is visible by eye under a fluorescence dissection scope, and can be recorded with short exposure time. No signal is observed when only the vector (B) or the target locus (C, D) is linearized, as also shown in Figure 2B. In the absence of exposed arms of homology, no signal is observed (panels D and J, with the exception of two cells in one of the two embryos shown here for condition D). Conditions in which only the left arm of homology (E, F, G) is exposed yield a few mNG-positive cells, but at a frequency that is much lower than in conditions A and A’. Conditions in which only the right arm is exposed (H, I) yield even fewer positive cells.

**Supplementary Figure 4.**
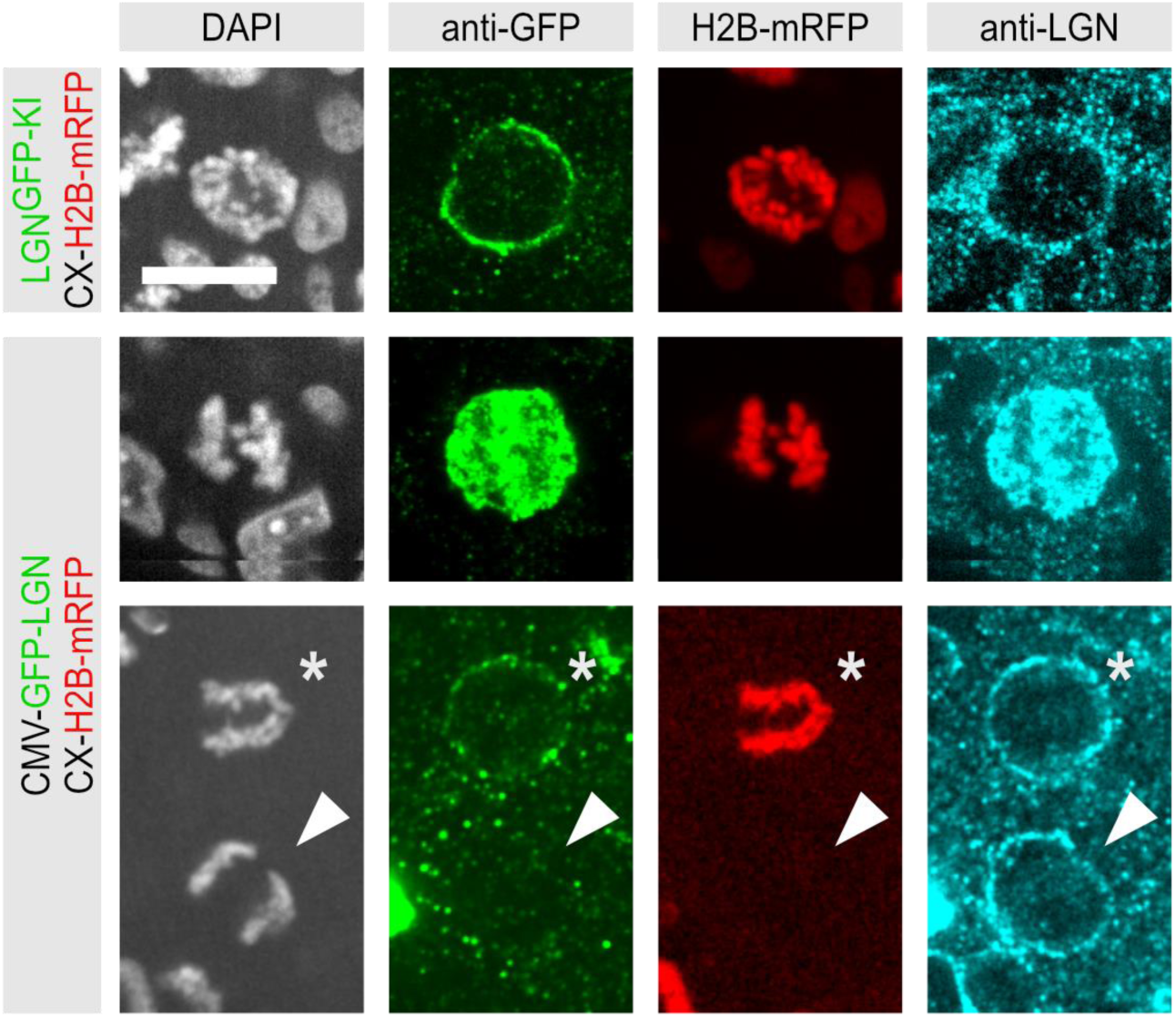
– KI reports subcellular distribution more faithfully than overexpression. A comparison of the subcellular distribution of GFP-tagged LGN (green) under knock-in (top row) and CMV-driven overexpression (middle and bottom rows), illustrated in early anaphase cells, and revealed with an anti-GFP antibody in en-face views of the fixed neuroepithelium. Endogenous and overexpressed LGN are also visualized with an anti-LGN antibody (cyan). An example of a wild type cell is provided for comparison (bottom row, white arrowhead). Expression of GFP-LGN under control of the CMV promoter results in either strong overexpression leading to saturating signal that fills the cytoplasm (middle row), or in low expression that results in a very low cortical GFP signal that recapitulates the endogenous LGN localization, but is barely detectable with the anti-GFP antibody (bottom row, asterisk). By comparison, the endogenously tagged LGN-GFP protein offers a better cortical to cytoplasmic signal ratio (top row). Scale bar, 10µm.

**Supplementary Figure 5.**
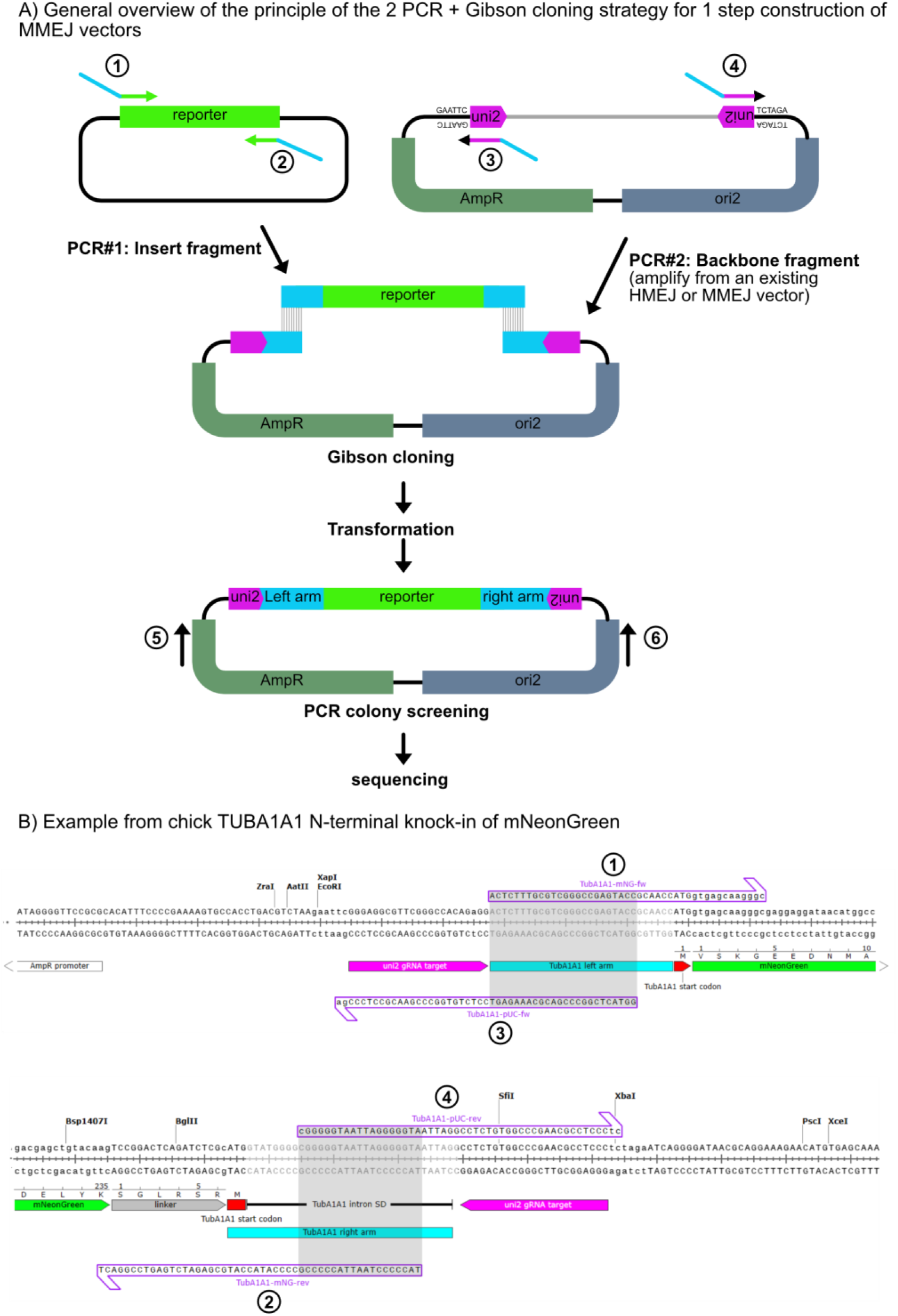
– overview of the single step cloning of MMEJ vetors. A. General overview of the principle: 2 PCR fragments spanning the reporter sequence and the vector backbone are generated with oligos including 5’ sequences designed to create ∼35bp arms of homology (blue) after Gibson recombination. B. Example from chick TUBA1A1 N-terminal knock-in of mNeonGreen: sequence and position of oligos 1, 2, 3 and 4 used for PCR, and sequence of junctions in final MMEJ vector after Gibson cloning. Gray boxes show the overlap sequence used for Gibson cloning.

**Supplementary Table 1.**
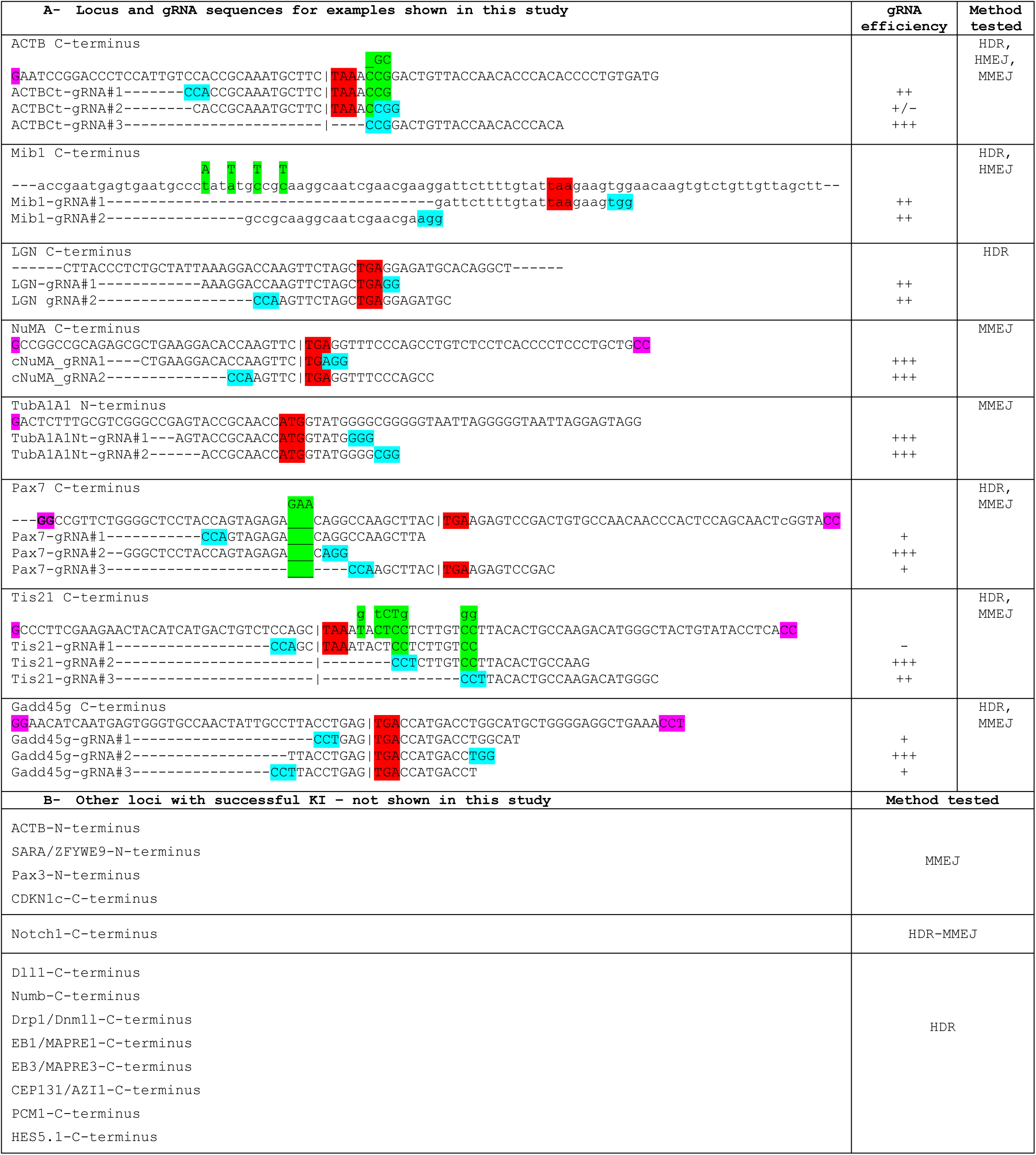
A. Locus and gRNA sequences for all genes targeted in this study. For each gene, the top line provides the genomic sequence surrounding the Start (N-terminal fusions) or Stop (C-terminal fusions) regions. The other lines show the sequence targeted by the different gRNAs that have been tested for each locus. ATG and stop codons are highlighted in red. Sequences highlighted in turquoise correspond to the gRNA target PAM sequences. Sequences highlighted in magenta show the 5’ and 3’ ends of short arms of homology used in MMEJ vectors which also partially match with the PAM in the donor vector linearization target site. Sequences highlighted in green over the ACTB, Mib1, Tis21 and Pax7 genomic sequence correspond to bases that have been changed/added/removed in the arms of homology of the donor vector to prevent targeting by gRNAs whose sequence would otherwise be entirely included in one arm of homology. The second column indicates the observed relative efficiency of each gRNA The third column indicates which donor vector architectures have been used for each locus B. A list of 13 other loci that have been targeted successfully, with the corresponding vector architecture

